# Genotype-by-environment interactions inferred from genetic effects on phenotypic variability in the UK Biobank

**DOI:** 10.1101/519538

**Authors:** Huanwei Wang, Futao Zhang, Jian Zeng, Yang Wu, Kathryn E. Kemper, Angli Xue, Min Zhang, Joseph E. Powell, Michael E. Goddard, Naomi R. Wray, Peter M. Visscher, Allan F. McRae, Jian Yang

## Abstract

Genotype-by-environment interaction (GEI) is a fundamental component in understanding complex trait variation. However, it remains challenging to identify genetic variants with GEI effects in humans largely because of the small effect sizes and the difficulty of monitoring environmental fluctuations. Here, we demonstrate that GEI can be inferred from genetic variants associated with phenotypic variability in a large sample without the need of measuring environmental factors. We performed a genome-wide variance quantitative trait locus (vQTL) analysis of ~5.6 million variants on 348,501 unrelated individuals of European ancestry for 13 quantitative traits in the UK Biobank, and identified 75 significant vQTLs with *P*<2.0×10^−9^ for 9 traits, especially for those related to obesity. Direct GEI analysis with five environmental factors showed that the vQTLs were strongly enriched with GEI effects. Our results indicate pervasive GEI effects for obesity-related traits and demonstrate the detection of GEI without environmental data.

## Introduction

Most human traits are complex because they are affected by many genetic and environmental factors as well as potential interactions between them^1,2^. Despite the long history of effort^3–5^, there has been limited success in identifying genotype-by-environment interaction (GEI) effects in humans^5–8^. This is likely because many environmental exposures are unknown or difficult to record during the life course, and because the effect sizes of GEI are small given the polygenic nature of most human traits^9–11^ so that the sample sizes of most previous studies are not large enough to detect the small GEI effects.

The GEI effect of a genetic variant on a quantitative trait could lead to differences in variance of the trait among groups of individuals with different variant genotypes (Figure 1a-b). GEI effects can therefore be inferred from a variance quantitative trait locus (vQTL) analysis^12^. Unlike the classical quantitative trait locus (QTL) analysis that tests the allelic substitution effect of a variant on the mean of a phenotype (Figure 1c), vQTL analysis tests the allelic substitution effect on the trait variance (Figure 1b or 1d). In comparison to the analyses that perform direct GEI tests, vQTL analysis could be a more powerful approach to identify GEI because it does not require measures of environmental factors and thus can be performed in data with very large sample sizes^13^. Although there had been empirical evidence for the genetic control of phenotypic variance in livestock for decades^14,15^, it was not until recent years that genome-wide vQTL analysis was applied in humans^12,16,17^, and only a handful of vQTLs have been identified for a limited number of traits (e.g. the *FTO* locus for body mass index (BMI)^17^) owing to the small effect sizes of the vQTLs. The availability of data from large biobank-based genome-wide association studies (GWAS)^18,19^ provide an opportunity to interrogate the genome for vQTLs for a range of phenotypes in cohorts with unprecedented sample size.

**Figure 1.**
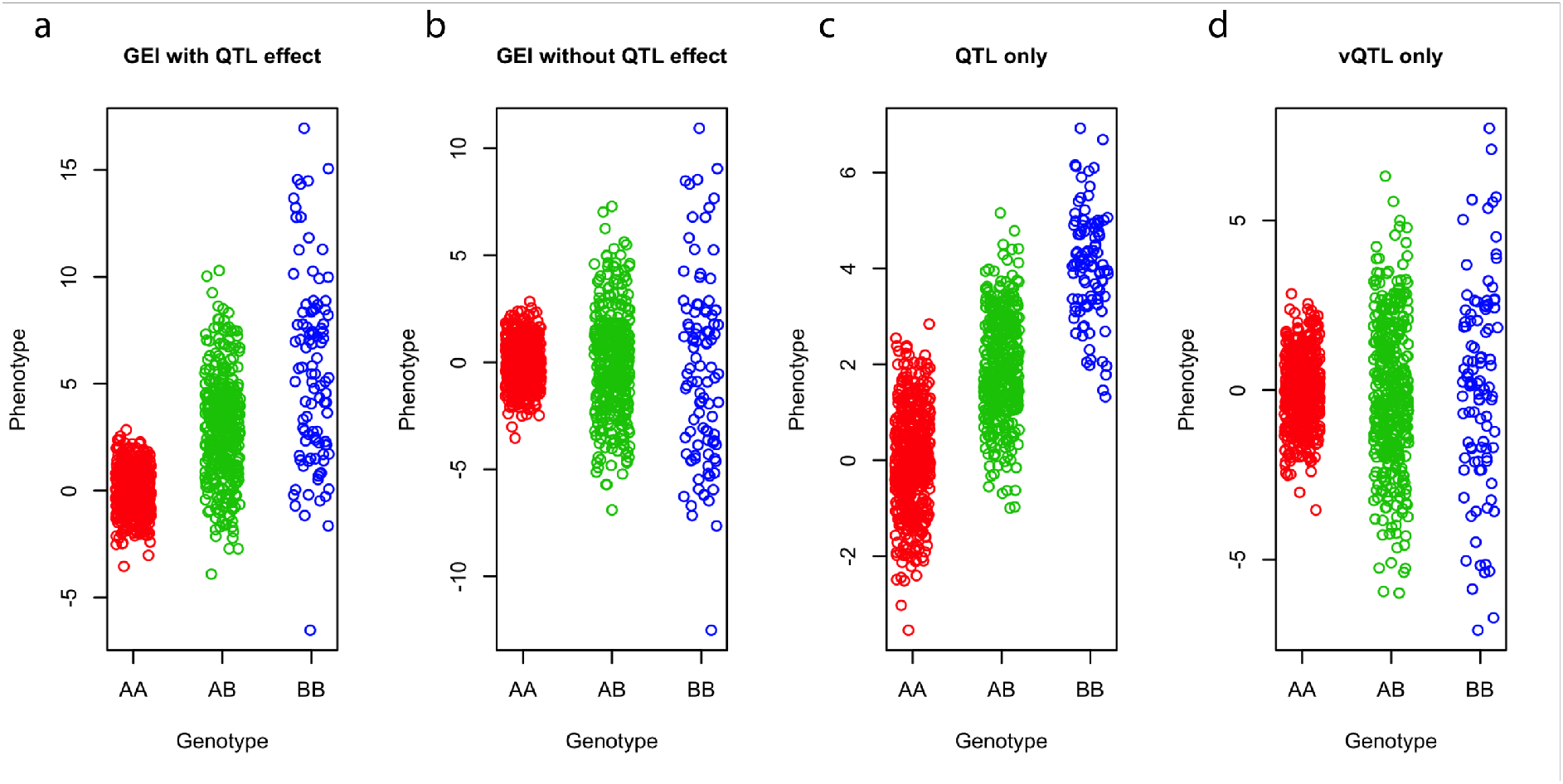
Schematic of the differences in mean or variance among genotype groups in the presence of GEI, QTL and vQTL effect. The phenotypes of 1,000 individuals were simulated based on a genetic variant (MAF = 0.3) with a) both QTL and GEI effects, (b) GEI effect only (no QTL effect), (c) QTL effect only (no GEI or vQTL effect), or (d) vQTL only (no QTL effect).

On the other hand, the statistical methods for vQTL analysis are not entirely mature^13^. There have been a series of classical non-parametric methods^20^, originally developed to detect violation of the homogeneous variance assumption in linear regression model, which can be used to detect vQTLs, including the Bartlett’s test^21^, the Levene’s test^22,23^ and the Fligner-Killen test^24^. Recently, more flexible parametric models have been proposed, including the double generalized linear model (DGLM)^25–27^ and the likelihood ratio test^28^. In addition, it has been suggested that the transformation of phenotype that alters phenotype distribution also has an influence on the power and/or false positive rate (FPR) of a vQTL analysis^16,29^.

In this study, we calibrated the most commonly used statistical methods for vQTL analysis by extensive simulations. We then used the best performing method to conduct a genome-wide vQTL analysis for 13 quantitative traits in 348,501 unrelated individuals using the full release of the UK Biobank (UKB) data^18^. We further investigated whether the detected vQTLs are enriched for GEI by conducting a direct GEI test for the vQTLs with five environmental factors.

## Results

### Evaluation of the vQTL methods by simulation

We used simulations to quantify the FPR and power (i.e., true positive rate) for the vQTL methods and phenotype processing strategies (Methods). We first simulated a quantitative trait based on a simulated single nucleotide polymorphism (SNP), i.e., a single-SNP model, under a number of different scenarios, namely: 1) five different distributions for the random error term (i.e., individual-specific environment effect); 2) four different types of SNP with or without the effect on mean or variance (Methods). We used the simulated data to compare four most widely used vQTL methods, namely the Bartlett’s test^21^, the Levene’s test^22,23^, the Fligner-Killen (FK) test^24^ and the DGLM^25–27^. We observed no inflation in FPR for the Levene’s test under the null (i.e., no vQTL effect) regardless of the skewness or kurtosis of the phenotype distribution or the presence or absence of the SNP effect on mean (Supplementary Figure 1a). These findings are in line with the results from previous studies^16,20,30^ that demonstrate the Levene’s test is robust to the distribution of phenotype. The FPR of the Bartlett’s test or DGLM was inflated if the phenotype distribution was skewed or heavy-tailed (Supplementary Figure 1a). The FK test seemed to be robust to kurtosis but vulnerable to skewness of the phenotype distribution (Supplementary Figure 1a). We also observed that logarithm or rank-based inverse-normal transformation (RINT) could result in inflated test statistics in the presence of QTL effect (i.e., SNP effect on mean; Supplementary Figure 1b).

To simulate more complex scenarios, we used a multiple-SNP model with two covariates (age and sex) and different numbers of SNPs (Figure 2). The results were similar to those observed above, although the power of the Levene’s test decreased with an increase of the number of causal SNPs (Figure 2a). Again, logarithm transformation or RINT gave rise to an inflated FPR in the presence of SNP effect on mean, and RINT led to a further loss of power (Figure 2b). These results also suggested that pre-adjusting the phenotype by covariates slightly increased the power of vQTL detection (Figure 2b). We therefore used the Levene’s test for real data analysis with the phenotypes pre-adjusted for covariates without logarithm transformation or RINT.

**Figure 2.**
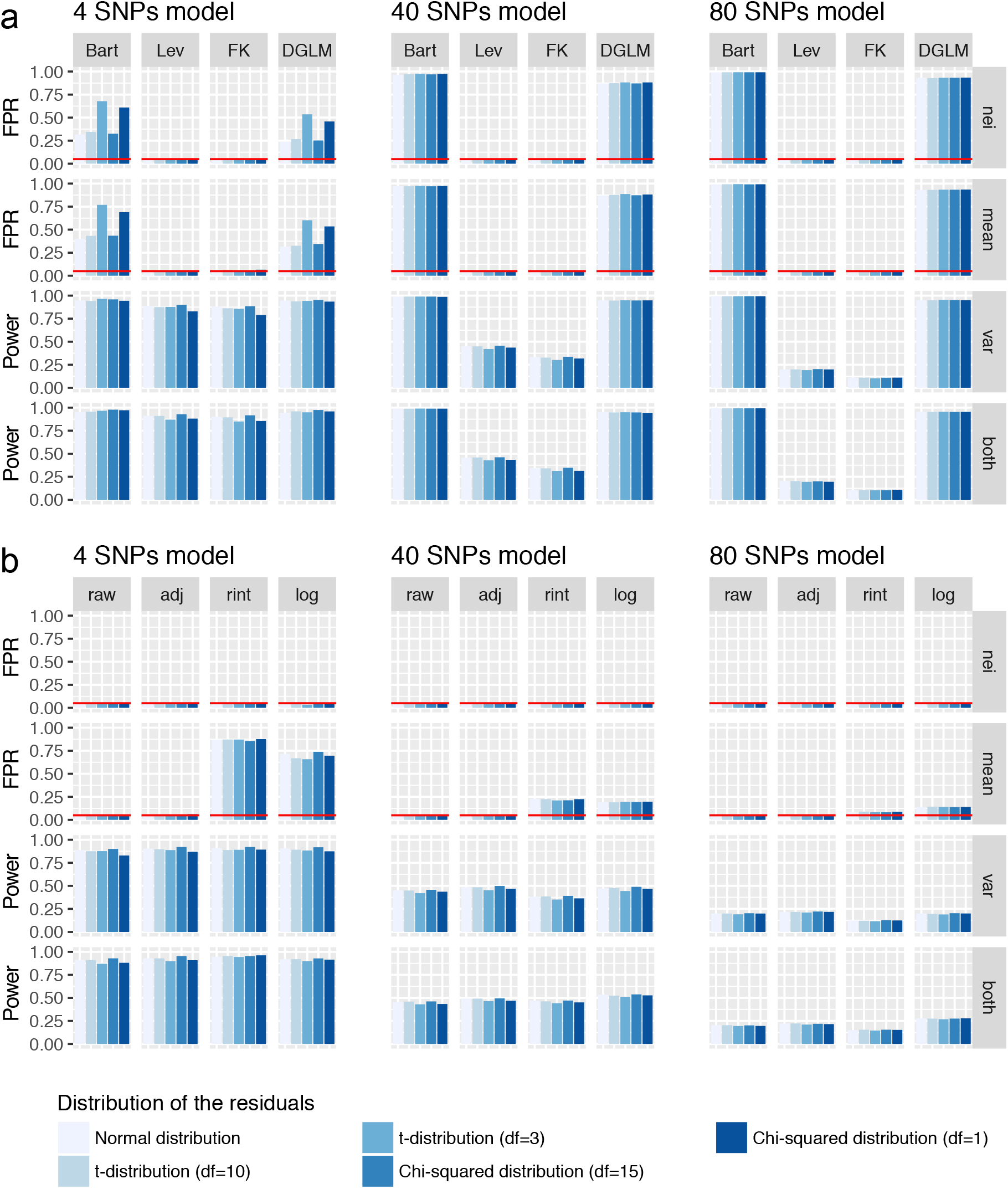
Evaluation of the statistical methods and phenotype processing strategies for vQTL analysis by simulation. Phenotypes of 10,000 individuals were simulated based on different number of SNPs (i.e. 4, 40 or 80), two covariates (i.e. sex and age) and one error term in a multiple-SNP model (Methods). The SNP effects were simulated under four scenarios: 1) effect on neither mean nor variance (nei), 2) effect on mean only (mean), 3) effect on variance only (var), or 4) effect on both mean and variance (both). The error term was generated from five different distributions: normal distribution, t-distribution with df = 10 or 3, or *χ^2^* distribution with df = 15 or 1. In panel a, four statistical test methods, i.e., the Bartlett’s test (Bart), the Levene’s test (Lev), the Fligner-Killen test (FK) and the DGLM, were used to detect vQTLs. In panel b, the Levene’s test was used to analyse phenotypes processed using four strategies, i.e., raw phenotype (raw), raw phenotype adjusted for covariates (adj), rank-based inverse-normal transformation after covariate adjustment (rint), and logarithm transformation after covariate adjustment (log). The FPR or power was calculated as the number of vQTLs with p < 0.05 divided by the total number of tests across 1,000 simulations. The red horizontal line represents an FPR of 0.05.

### Genome-wide vQTL analysis for 13 UKB traits

We performed a genome-wide vQTL analysis using the Levene’s test with 5,554,549 genotyped or imputed common variants on 348,501 unrelated individuals of European ancestry for 13 quantitative traits in the UKB^18^ (Methods, Supplementary Table 1 and Supplementary Figure 2). For each trait, we pre-adjusted the phenotype for age and the first 10 principal components (PCs, derived from SNP data) and standardised the residuals to z-scores in each gender group (Methods). This process removed not only the effects of age and the first 10 PCs on the phenotype but also the differences in mean and variance between the two genders. We excluded individuals with adjusted phenotypes more than 5 standard deviations (SD) from the mean and removed SNPs with minor allele frequency (MAF) smaller than 0.05 to avoid potential false positive associations due to the coincidence of a low-frequency variant with an outlier phenotype (see Supplementary Figure 3 for an example). We acknowledge that this process could potentially result in a loss of power, but this can be compensated for by the use of a very large sample (n ~ 350,000).

With an experiment-wise significant threshold 2.0×10^−9^ (i.e., 1×10^−8^/5.03 with 1×10^−8^ being a more stringent genome-wide significant threshold recommended by recent studies^31,32^ and 5.03 being the effective number of independent traits (Supplementary Note 3)), we identified 75 vQTLs for 9 traits (Figure 3, Table 1 and Supplementary Table 2). There was no vQTL for height, consistent with the observation in a previous study^17^. We identified more than 15 vQTLs for each of the three obesity-related traits, i.e., BMI, waist circumference (WC), and hip circumference (HC) (Table 1). The 75 vQTLs were located at 40 near-independent loci after excluding one of each pair of top vQTL SNPs (i.e., the SNP with lowest vQTL p-value at each vQTL association peak) with linkage disequilibrium (LD) r^2^ > 0.01, suggesting that some of the loci were associated with the phenotypic variance of multiple traits. For example, the *FTO* locus was associated with the phenotypic variance of WC, HC, BMI, body fat percentage (BFP) and basal metabolic rate (BMR) (Figure 4). For the lung-function-related traits, there was no significant vQTL for forced expiratory volume in one second (FEV1) and forced vital capacity (FVC) but were 3 vQTLs for FEV1/FVC ratio (FFR).

**Figure 3.**
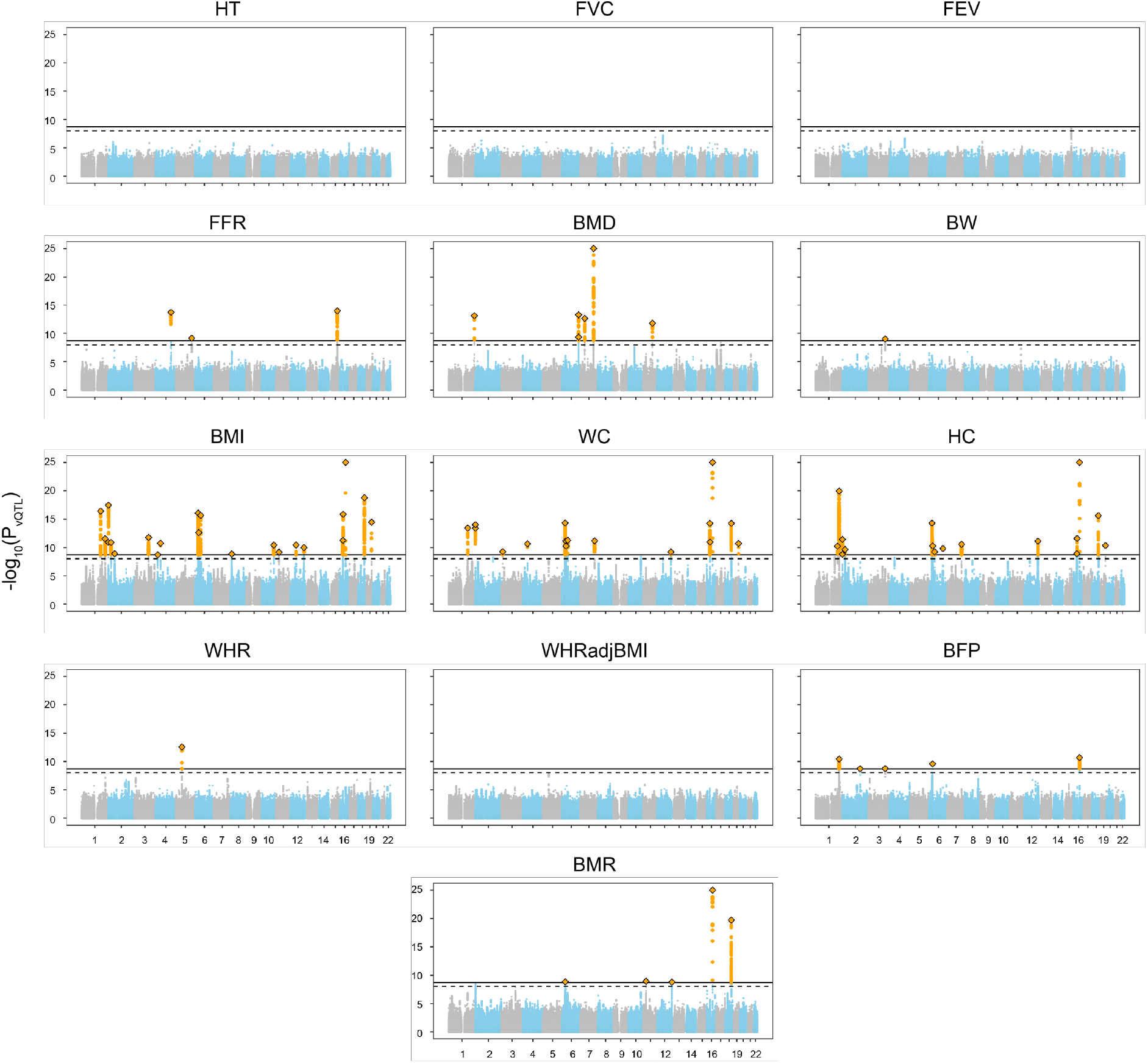
Manhattan plots of genome-wide vQTL analysis for 13 traits in the UKB. For each of the 13 traits (see Table 1 for full names of the traits), test statistics (−log_10_(*P*_vQTL_)) of all common (MAF≥0.05) SNPs from the vQTL analysis are plotted against their physical positions. The dash line represents the genome-wide significance level 1.0 ×10^−8^ and the solid line represents the experiment-wise significance level 2.0×10^−9^. For graphical clarity, SNPs with *P*_vQTL_ < 1×10^−25^ are omitted, SNPs with *P*_vQTL_ < 2.0×10^−9^ are colour-coded in orange, the top vQTL SNP is represented by a diamond, and the remaining SNPs are colour-coded in grey or blue for odd or even chromosome.

**Table 1.** The number of experiment-wise significant vQTLs or QTLs for the 13 UKB traits.

| Trait | Description | Number<br>of vQTLs | Number of<br>QTLs |
| --- | --- | --- | --- |
| HT | Standing height | 0 | 1063 |
| FVC | Forced vital capacity | 0 | 325 |
| FEV1 | Forced expiratory volume in 1-second | 0 | 266 |
| FFR | FEV1 and FVC ratio | 3 | 17 |
| BMD | Heel bone mineral density T-score, automated | 6 | 267 |
| BW | Birth weight | 1 | 57 |
| BMI | Body mass index | 22 | 271 |
| WC | Waist circumference | 16 | 196 |
| HC | Hip circumference | 16 | 249 |
| WHR | Waist to Hip Ratio | 1 | 157 |
| WHRadjBMI | WHR adjusted for BMI | 0 | 221 |
| BFP | Body fat percentage | 5 | 249 |
| BMR | Basal metabolic rate | 5 | 465 |
| Total |  | 75 | 3,803 |

**Figure 4.**
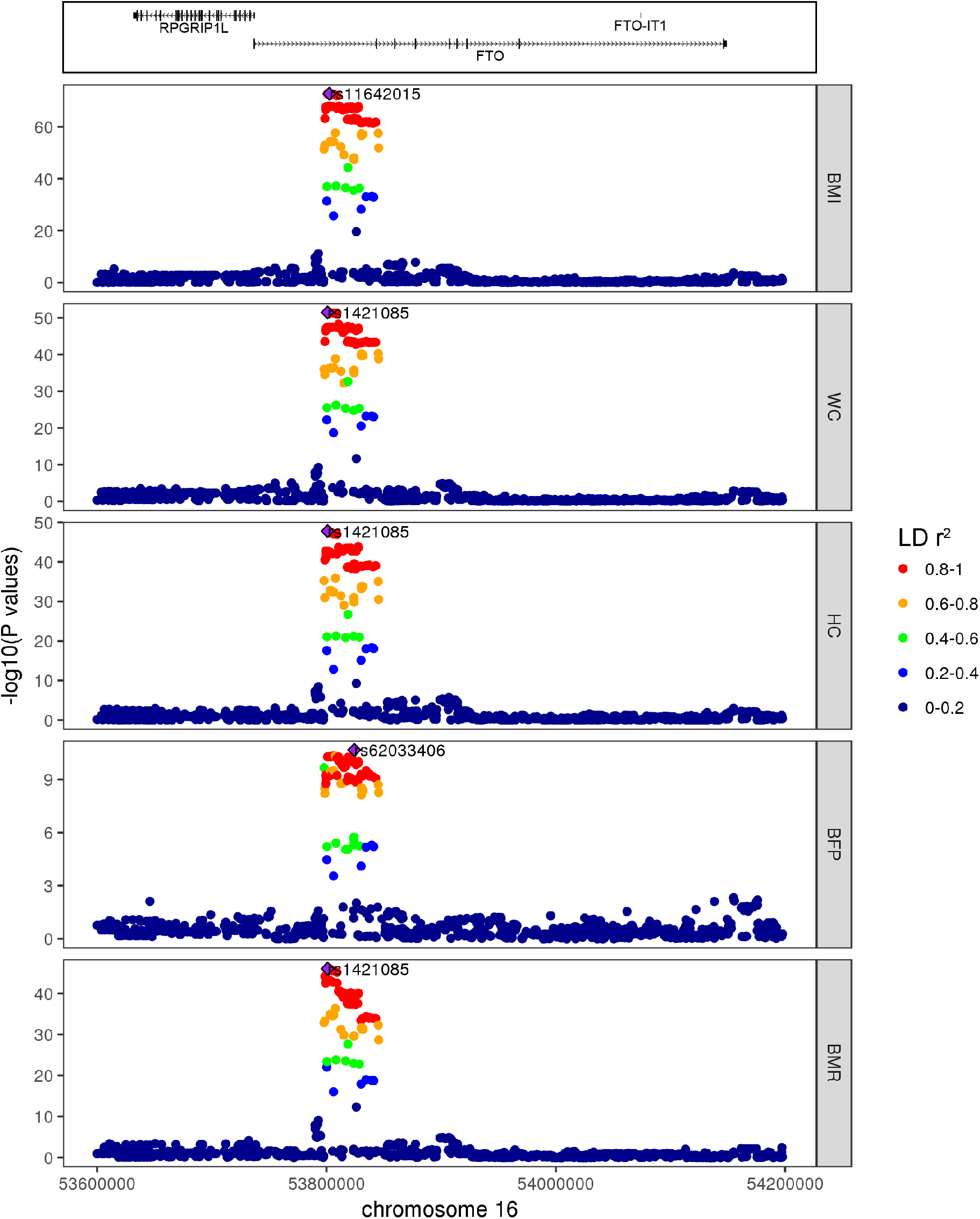
Regional plots of the *FTO* locus associated with the phenotypic variability of 5 traits. For each of these 5 traits for which the phenotypic variance is significantly associated with the *FTO* locus, vQTL test statistics (−log_10_(P_vQTL_)) are plotted against SNP positions surrounding the top vQTL SNP (represented by a purple diamond) at the *FTO* locus. SNPs in different levels of LD with the top vQTL SNP are shown in different colours. The RefSeq genes in the top panel are extracted from the UCSC Genome Browser (URLs).

The Levene’s test assesses the difference in variance among three genotype groups free of the assumption about additivity (i.e., the vQTL effect of carrying two copies of the effect allele is not assumed to be twice that carrying one copy). We found two vQTLs (i.e., rs141783576 and rs10456362) potentially showing non-additive genetic effect on the variance of HC and BMR, respectively (Supplementary Table 2).

### GWAS analysis for the 13 UKB traits

To investigate whether the SNPs with effects on variance also have effects on mean, we performed GWAS (or genome-wide QTL) analyses for the 13 UKB traits described above. We identified 3,803 QTLs at an experiment-wise significance level (i.e., *P*_QTL_ < 2.0×10^−9^) for the 13 traits in total, a much larger number than that of the vQTLs (Table 1 and Figure 5). Among the 75 vQTLs, the top vQTL SNPs at 9 loci did not pass the experiment-wise significance level in the QTL analysis (Supplementary Table 2). For example, the *CCDC92* locus showed a significant vQTL effect but no significant QTL effect on WC (Supplementary Table 2 and Figure 6a), whereas the *FTO* locus showed both significant QTL and vQTL effects on WC (Figure 6b). For the 66 vQTLs with both QTL and QTL effects, the vQTL effects were all in the same directions as the QTL effects, meaning that for any of these SNPs the genotype group with larger phenotypic mean also tends to have larger phenotypic variance than the other groups. For the 9 loci with vQTL effects only, it is equivalent to a scenario where a QTL has a GEI effect with no (or a substantially reduced) effect on average across different levels of an environmental factor (Figure 1b).

**Figure 5.**
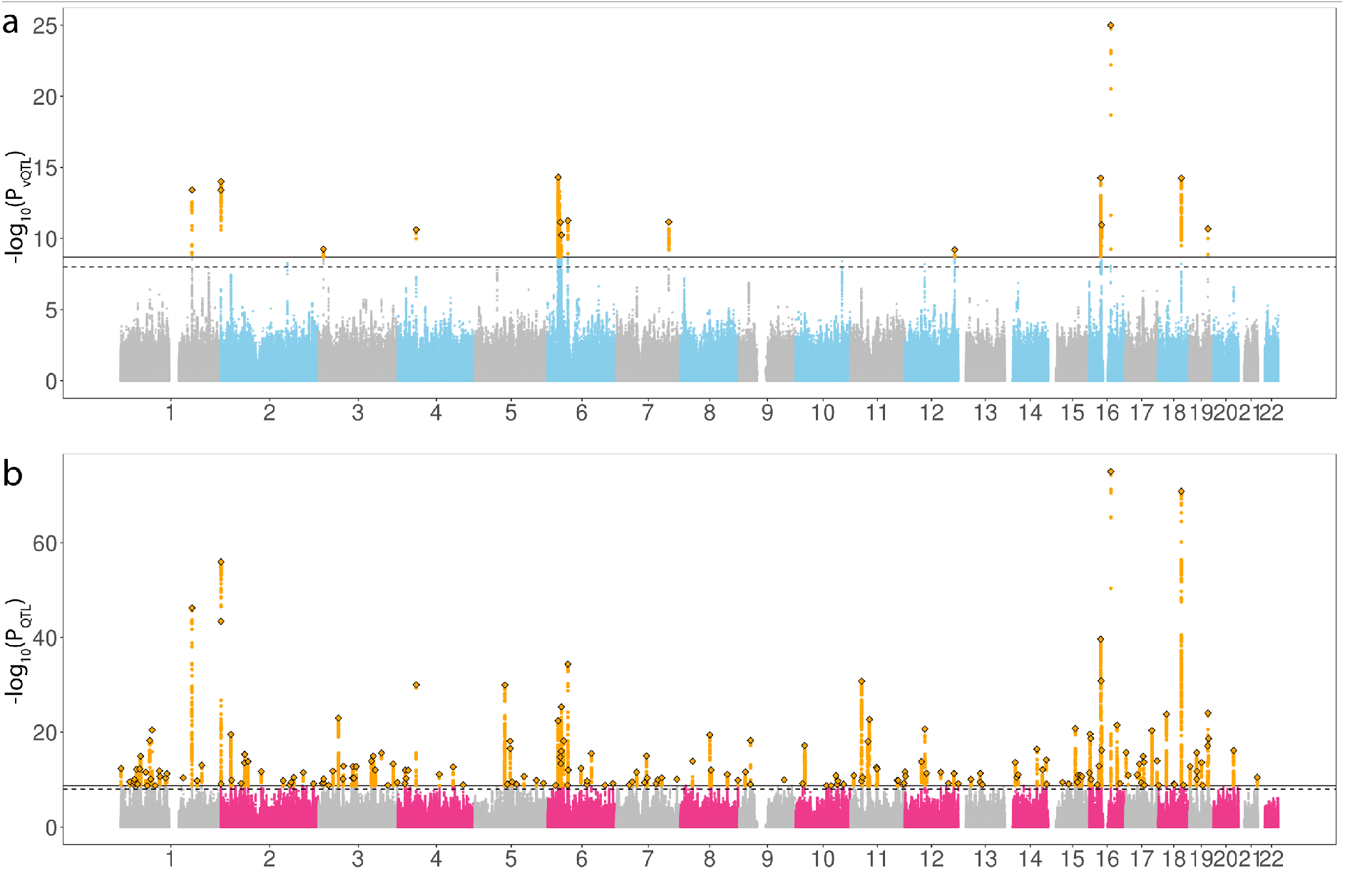
Manhattan plots of genome-wide vQTL or QTL analysis for waist circumference in the UKB. Test statistics (−log_10_(*P*_vQTL_)) of all common SNPs from vQTL (a) or QTL (b) analysis are plotted against their physical positions. The dash line represents the genome-wide significance level 1×10^−8^ and the solid line represents the experiment-wise significance level 2.0×10^−9^. For graphical clarity, SNPs with *P*_vQTL_ < 1×10^−25^ or *P*_QTL_ < 1×10^−75^ are omitted, SNPs with *P* < 2.0×10^−9^ are colour-coded in orange, the top vQTL or QTL SNP is represented by a diamond, and the remaining SNPs are colour-coded in grey or blue for vQTL analysis (a) or grey or pink for QTL analysis (b) for odd or even chromosomes.

**Figure 6.**
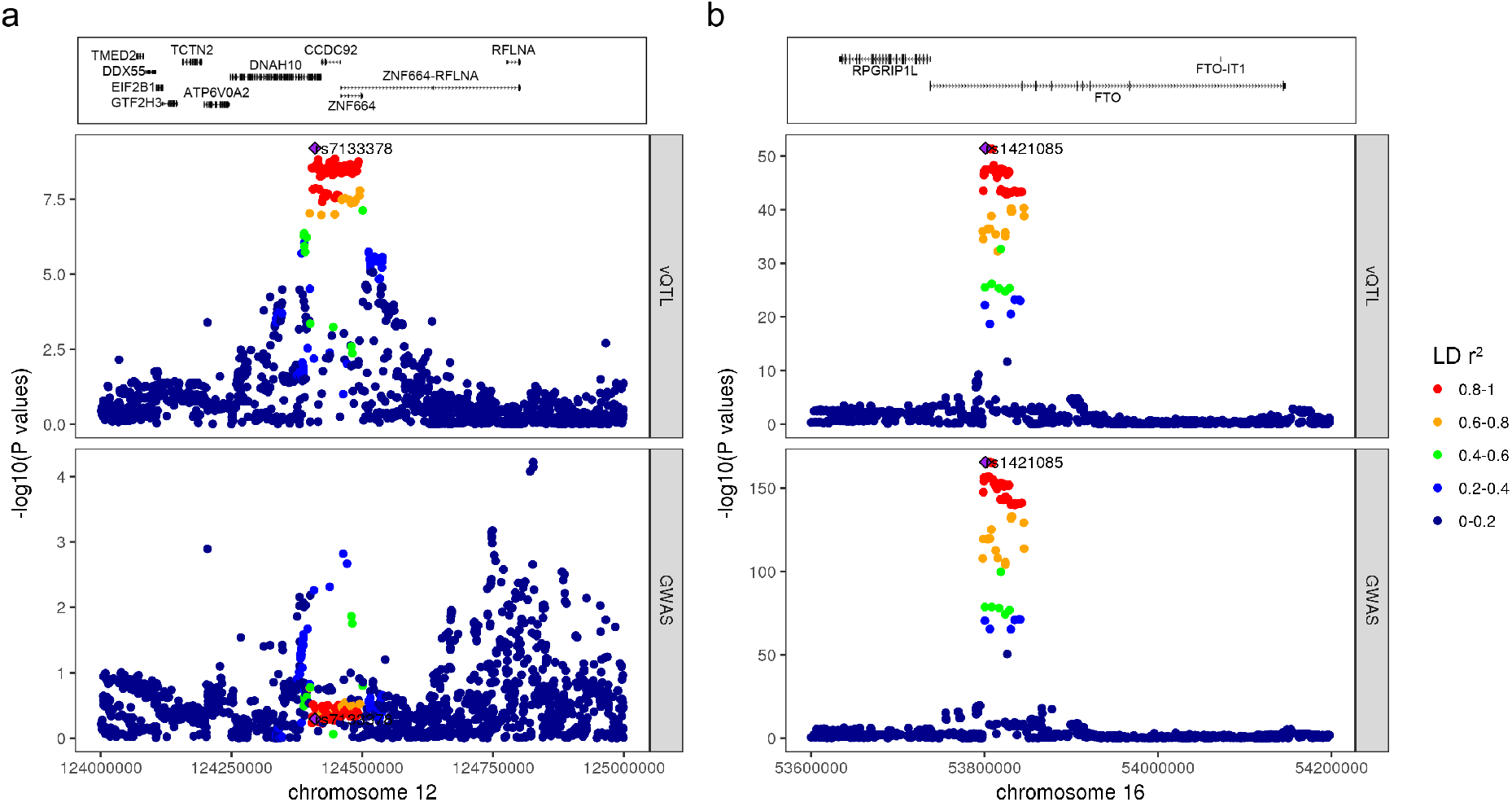
QTL and vQTL regional plots of the *CCDC92* or *FTO* locus for waist circumference. The QTL and vQTL test statistics (i.e., -log_10_(*P* values)) for waist circumference are plotted against SNP positions surrounding the top vQTL SNP at the *CCDC92* (panel a) or *FTO* locus (panel b). The top vQTL SNP is represented by a purple diamond. SNPs in different levels of LD with the top vQTL SNP are shown in different colours. The RefSeq genes in the top panel are extracted from the UCSC Genome Browser (URLs).

### vQTL and GEI

To further investigate whether the associations between vQTLs and phenotypic variance can be explained by GEI, we performed a direct GEI test based on an additive genetic model with an interaction term between a top vQTL SNP and one of five environmental/covariate factors in the UKB data (Methods). The five environmental factors are sex, age, physical activity (PA), sedentary behaviour (SB), and ever smoking (Supplementary Note 4, Supplementary Figure 4 and Supplementary Table 3). We observed 16 vQTLs showing a significant GEI effect with at least one of five environmental factors after correcting for multiple tests (p < 1.3×10^−4^ = 0.05/(75*5); Figure 7a and Supplementary Table 4).

**Figure 7.**
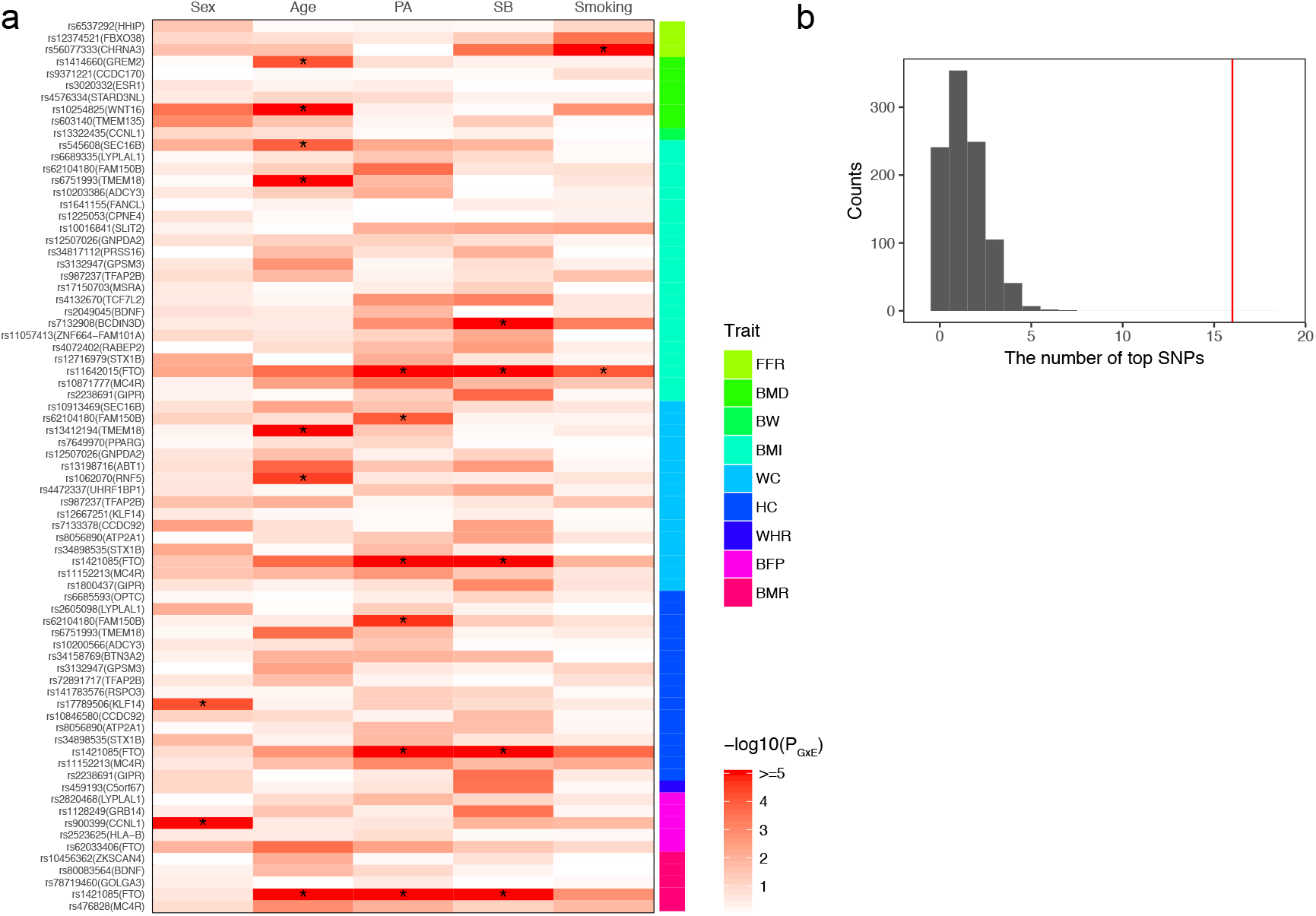
Enrichment of GEI effects among the 75 vQTLs in compared with a random set of QTLs. Five environmental factors, i.e., sex, age, physical activity (PA), sedentary behaviour (SB), and smoking, were used in the GEI analysis. (a) The heatmap plot of GEI test statistics (−log_10_(P_GEI_)) for the 75 top vQTL SNPs. “*” denotes significant GEI effects after Bonferroni correction (P_GEI_ < 1.33×10^−4^ = 0.05/(75*5)). (b) The distribution of the number of significant GEI effects for 75 top QTL SNPs randomly selected from all the top QTL SNPs with 1000 repeats (mean 1.39 and SD 1.15). The red line represents the number of significant GEI effects for the 75 top vQTL SNPs (i.e., 16).

To test whether the GEI effects are enriched among vQTLs in comparison with the same number of QTLs, we performed GEI test for 75 top GWAS SNPs randomly selected from all the QTLs and repeated the analysis 1000 times. Of the 75 top SNPs with QTL effects, the number of SNPs with significant GEI effects was 1.39 averaged from the 1000 repeated samplings with a SD of 1.15 (Figure 7b), significantly lower the number (16) observed for the vQTLs (the difference is larger than 12 SDs, equivalent to p = 6.6×10^−37^). This result shows that SNPs with vQTL effects are much more enriched with GEI effects compared to those with QTL effects. To exclude the possibility that the GEI signals were driven by phenotype processing (e.g., the adjustment of phenotype for sex and age), we repeated the GEI analyses using raw phenotype data without covariates adjustment; the results remain largely unchanged (Supplementary Figure 5).

## Discussion

In this study, we leveraged the genetic effects associated with phenotypic variability to infer GEI. We calibrated the most commonly used vQTL methods by simulation. We found that the FPR of the Levene’s test was well-calibrated across all simulation scenarios whereas the other methods showed an inflated FPR if the phenotype distribution was skewed or heavy-tailed under the null hypothesis (i.e., no vQTL effect), despite that the Levene’s test appeared to be less powerful than the other methods under the alternative hypothesis in particular when the per-variant vQTL effect was small (Figure 2 and Supplementary Figure 1). Parametric bootstrap or permutation procedures have been proposed to reduce the inflation in the test-statistics of DGLM and LRT-based method, both of which are expected to be more powerful than the Levene’s test^28,30^, but bootstrap and permutation are computationally inefficient and thus not practically applicable to biobank data such as the UKB. In addition, we observed inflated FPR for the Levene’s test in the absence of vQTL effects but in the presence of QTL effects if the phenotype was transformed by logarithm transformation or RINT. We therefore recommend the use of the Levene’s test in practice without logarithm transformation or RINT of the phenotype. In addition, a very recent study by Young et al.^33^ developed an efficient algorithm to perform a DGLM analysis and proposed a method (called dispersion effect test (DET)) to remove the founding in vQTL associations (identified by DGLM) due to the QTL effects. We showed by simulation that when the number of simulated causal variants was relatively large (note that the DET test is not applicable to oligogenic traits), the Young et al. method (DGLM followed by DET) performed similarly as the Levene’s test with differences depending on how the phenotype was processed (Supplementary Figure 6).

We identified 75 genetic variants with vQTL effects for 9 quantitative traits in the UKB at a stringent significance level and observed strong enrichment of GEI effects among the genetic variants with vQTL effects compared to those with QTL effects. There are several vQTLs for which the GEI effect has been reported in previous studies. The first example is the interaction effect of the *CHRNA5-A3-B4* locus (rs56077333) with smoking lung function (as measured by FFR ratio, i.e., FEV1/FVC), *P*_vQTL_ = 1.1×10^−14^ and *P*_GEI(smoking)_ = 4.6×10^−25^ (Supplementary Table 2 and 4). The *CHRNA5-A3-B4* gene cluster is known to be associated with smoking and nicotine dependence^34–36^. However, results from recent GWAS studies^37–39^ do not support the association of this locus with lung function. We hypothesize that the effect of the *CHRNA5-A3-B4* locus on lung function depends on smoking^40^ (Supplementary Table 5). The vQTL signal at this locus remained (*P*_vQTL_ = 5.2×10^−12^) after adjusting the phenotype for array effect, which was reported to affect the QTL association signal at this locus^18^. The second example is the interaction of the *WNT16-CPED1* locus with age for BMD (rs10254825: *P*_vQTL_ = 2.0×10^−45^ and PGEI(age) = 1.2×10^−7^). The *WNT16-CPED1* locus is one of the strongest BMD-associated loci identified from GWAS^41,42^.

We observed a genotype-by-age interaction effect at this locus for BMD (Supplementary Table 6), in line with the results from previous studies that the effect of the top SNP at *WNT16-CPED1* on BMD in humans^43^ and the knock-out effect of *Wnt16* on bone mass in mice^44^ are age-dependent. The third example is the interaction of the *FTO* locus with physical activity and sedentary behaviour for obesity-related traits (*P*_vQTL_ < 1×10^−10^ for BMI, WC, HC, BFP and BMR; PGEI(PA) = 1.3×10^−10^ for BMI, 1.4×10^−7^ for WC, 5.3×10^−7^ for HC and 2.6×10^−7^ for BMR). The *FTO* locus was one of the first loci identified by the GWAS of obesity-related traits^45^ although subsequent studies^46,47^ show that *IRX3* and *IRX5* (rather than *FTO)* are the functional genes responsible for the GWAS association. The top associated SNP at the *FTO* locus is not associated with physical activity but its effect on BMI decreases with the increase of physical activity level^48,49^, consistent with the interaction effects of the *FTO* locus with physical activity or sedentary behaviour for obesity-related traits identified in this study (Supplementary Tables 7 and 8). In addition, 5 of the 22 BMI vQTLs were in LD (r^2^ > 0.5) with the variants (identified by a recently developed multiple-environment GEI test) showing significant interaction effects at FDR < 5% (corresponding to p < 1.16×10^−3^) with at least one of 64 environmental factors for BMI in the UKB^50^.

Apart from GEI, there are other possible interpretations of an observed vQTL signal, including “phantom vQTLs”^28,51^ and epistasis (genotype-by-genotype interaction). If the underlying causal QTL is not well imputed or not well tagged by a genotyped/imputed variant, the untagged variation at the causal QTL will inflate the vQTL test-statistic, potentially leading to a spurious vQTL association, i.e., the so-called phantom vQTL. We showed by theoretical deviations that the Levene’s test-statistic due to the phantom vQTL effect was a function of sample size, effect size of the causal QTL, allele frequency of the causal QTL, allele frequency of the phantom vQTL, and LD between the causal QTL and the phantom vQTL (Supplementary Note 5 and Supplementary Figure 7). From our deviations, we computed the numerical distribution of the expected phantom vQTL F-statistics given a number of parameters including the sample size (n = 350,000), variance explained by the causal QTL (*q*^2^ = 0.005, 0.01 or 0.02), and MAFs of the causal QTL and the phantom vQTL (MAF = 0.05 – 0.5). The result showed that for a causal QTL with q^2^ < 0.005 and MAF > 0.05, the largest possible phantom vQTL F-statistic was smaller than 2.69 (corresponding to a p-value of 6.8×10^−2^; Supplementary Figure 8). This explains why there were thousands of genome-wide significant QTLs but no significant vQTL for height (Table 1 and Figure 3). This result also suggests that the vQTLs detected in this study are very unlikely to be phantom vQTLs because the estimated variance explained by their QTL effects were all smaller than 0.005 except for rs10254825 at the *WNT16* locus on BMD (q^2^ = 0.014) (Supplementary Figure 9). However, our numerical calculation also indicated that for a QTL with MAF > 0.3 and *q*^2^ < 0.02, the largest possible phantom vQTL F-statistic was smaller than 5.64 (corresponding to a p-value of 3.6×10^−3^), suggesting rs10254825 is also unlikely to be a phantom vQTL. Note that we used the variance explained estimated at the top GWAS SNP to approximate q^2^ of the causal QTL so that q^2^ was likely to be underestimated because of imperfect tagging. However, considering the extremely high imputation accuracy for common variants^52^, the strong LD between the causal QTLs and the GWAS top SNPs observed in a previous simulation study based on whole-genome-sequence data^31^, and the overestimation of variance explained by the GWAS top SNPs because of winner’s curse, the underestimation in causal QTL q^2^ is likely to be small. In addition, we re-ran the vQTL analysis with the phenotype adjusted for the top GWAS variants within 10Mb distance of the top vQTL SNP; the vQTL signals after this adjustment were highly concordant with those without adjustment (Supplementary Figure 10). We further showed that there was no evidence for epistatic interactions between the top vQTL SNPs and any other SNP in more than 10 Mb distance or on a different chromosome (Supplementary Figure 11).

In conclusion, we systematically quantified the FPR and the power of four commonly used vQTL methods by extensive simulations and demonstrated the robustness of the Levene’s test. We also showed that in the presence of QTL effects the Levene’s test statistic could be inflated if the phenotype was transformed by logarithm transformation or RINT. We implemented the Levene’s test as part of the OSCA software package^53^ (URLs) for efficient genome-wide vQTL analysis, and applied OSCA-vQTL to 13 quantitative traits in the UKB and identified 75 vQTL (at 40 independent loci) associated with 9 traits, 9 of which did not show a significant QTL effect. As a proof-of-principle, we performed GEI analyses in the UKB with 5 environmental factors, and demonstrated the enrichment of GEI effects among the detected vQTLs. We further derived the theory to compute the expected “phantom vQTL” test-statistic due to untagged causal QTL effect, and showed by numerical calculation that our observed vQTLs were very unlikely to be driven by imperfectly tagged QTL effects. Our theory is also consistent with the observation of pervasive phantom vQTLs for molecular traits with large-effect QTLs (e.g., DNA methylation^51^). However, the conclusions from this study may be only applicable to quantitative traits of polygenic architecture. We caution vQTL analysis for binary or categorical traits, or molecular traits (e.g., gene expression or DNA methylation), for which the methods need further investigation.

## Methods

### Simulation study

We used a DGLM^25–27^ to simulate the phenotype based on two models with simulated SNP data in a sample of 10,000 individuals, i.e., a single-SNP model and multiple-SNP model with two covariates (i.e. age and sex). The single-SNP model can be written as

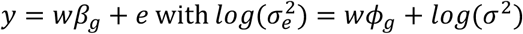

and the multiple-SNP model can be expressed as

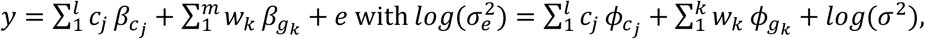

where *y* is a simulated phenotype; *w* or *w_k_* is a standardized SNP genotype, i.e., 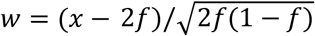 with *x* being the genotype indicator variable coded as 0, 1 or 2, generated from binomial(2, *f*) and *f* being the MAF generated from uniform(0.01, 0.5); *c_j_* is a standardized covariate with *c*_1_ (sex) generated from binomial(1, 0.5) and *C*_2_ (age) generated from uniform(20, 60); *e* is an error term normally distributed with mean 0 and variance 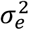. To simulate the error term with different levels of skewness and kurtosis, we generated *e* from five different distributions, including normal distribution, t-distribution with degree of freedom (df) = 10 or 3 and *χ^2^* distribution with df = 15 or 1. *β* (*φ*) is the effect on mean (variance) generated from *N*(0,1) if exists, 0 otherwise. *log*(*σ*^2^) is the intercept of the second linear model which was set to 0. We re-scaled the different components to control the variance explained, i.e., 0.1 and 0.9 for the genotype component and error term, respectively, for the single-SNP model, and 0.2, 0.4 and 0. 4 for the covariate component, genotype component and error term, respectively, for the multiple-SNP model. We simulated the SNP effects in four different scenarios: 1) effect on neither mean nor variance (nei), 2) effect on mean only (mean), 3) effect on variance only (var), or 4) effect on both mean and variance (both). We simulated only one causal SNP in the single-SNP model and 4, 40 or 80 causal SNPs in the multiple-SNP model.

We performed vQTL analyses using the simulated phenotype and SNP data to compare four vQTL methods, including the Bartlett’s test^21^, the Levene’s test^23^, the Fligner-Killeen test^24^ and the DGLM (Supplementary Note 1). We also performed the Levene’s test with four phenotype process strategies, including raw phenotype (raw), raw phenotype adjusted for covariates (adj), RNIT after covariate adjustment (rint), and logarithm transformation after covariate adjustment (log) (Supplementary Note 2). We repeated the simulation 1,000 times and calculated the FPR and power at p < 0.05 at a single SNP level.

### The UK Biobank data

The full release of the UKB data comprised of genotype and phenotype data for ~500,000 participates across the UK^18^. The genotype data were cleaned and imputed to the Haplotype Reference Consortium (HRC)^52^ and UK10K^54^ reference panels by the UKB team. Genotype probabilities from imputation were converted to hard-call genotypes using PLINK2^55^ (--hard-call 0.1). We excluded genetic variants with MAF < 0.05, Hardy-Weinberg equilibrium test p value < 1×10^−5^, missing genotype rate > 0.05 or imputation INFO score < 0.3, and retained 5,554,549 variants for analysis.

We identified a subset of individuals of European ancestry (n = 456,422) by projecting the UKB PCs onto those of 1000 Genome Project (1KGP)^56^. Furthermore, we removed one of each pair of individuals with SNP-derived (based on HapMap 3 SNPs) genomic relatedness > 0.05 using GCTA-GRM^57^ and retained 348,501 unrelated European individuals for further analysis.

We selected 13 quantitative traits for our analysis (Supplementary Table 1 and Supplementary Figure 2). The raw phenotype values were adjusted for age and the first 10 PCs in each gender group. We excluded from the analysis phenotype values that were more than 5 SD from the mean. The phenotypes were then standardized to z-scores with mean 0 and variance 1.

### Genome-wide vQTL analysis

The genome-wide vQTL analysis was conducted using the Levene’s test implemented in the software tool OSCA^53^ (URLs). The Levene’s test used in the study (also known as the median-based Levene’s test or the Brown-Forsythe test^23^) is a modified version of the original Levene’s test^22^ developed in 1960 that is essentially an one-way analysis of variance (ANOVA) of the variable 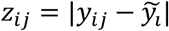, where *y_ij_* is phenotype of the *j-th* individual in the *i*-th group and 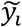 is the median of the *i*-th group. The Levene’s test statistic

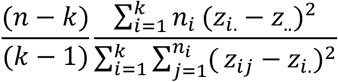

follows a *F* distribution with *k* – 1 and *n* – *k* degrees of freedom, where *n* is the total sample size, *k* is the number of groups (*k* = 3 in vQTL analysis), *n_i_* is the sample size of the *i*-th group,

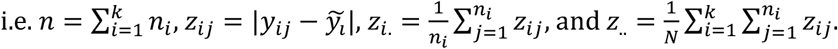

The experiment-wise significance level was set to 2.0×10^−9^, which is the genome-wide significance level (i.e. 1×10^−8^)^31,32^ divided by the effective number of independent traits (i.e. 5.03 for 13 traits). The effective number of independent traits was estimated based on the phenotypic correlation matrix^58^ (Supplementary Note 3). To determine the number of independent vQTLs, we performed an LD clumping analysis for each trait using PLINK2^55^ (--clump option with parameters --clump-p1 2.0e-9 --clump-p2 2.0e-9 --clump-r2 0.01 and -- clump-kb 5000). To visualize the results, we generated the Manhattan and regional association plots using ggplot2 package in R.

### GWAS analysis

The GWAS (or genome-wide QTL) analysis was conducted using PLINK2^55^ (--assoc option) using the same data as used in the vQTL analysis (note that the phenotype had been pre-adjusted for covariates and PCs). The other analyses, including LD clumping, and visualization, were performed using the same pipelines as those for genome-wide vQTL analysis described above.

### GEI analysis

Five environmental/covariate factors (i.e., sex, age, PA, SB and smoking) were used for the direct GEI tests. Sex was coded as 0 or 1 for female or male. Age was an integer number ranging from 40 to 74. PA was assessed by a three-level categorical score (i.e., low, intermediate and high) based on the short form of the International Physical Activity Questionnaire (IPAQ) guideline^59^. SB was an integer number defined as the combined time (hours) spent driving, non-work-related computer using or TV watching. The smoking factor “ever smoked” was coded as 0 or 1 for never or ever smoker. More details about the definition and derivation of environmental factor PA, SB and smoking can be found in the Supplementary Note 4, Figure 4 and Table 3.

We performed a GEI analysis to test the interaction effect between the top vQTL SNP and one of the five environmental factors based on the following model

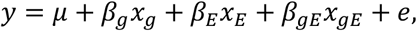

where *y* is phenotype, *μ* is the mean term, *x_g_* is mean-centred SNP genotype indicator, *x_E_* is mean-centred environmental factor, and *x_gE_ = x_g_x_E_*. We used a standard ANOVA analysis to test for *β_gE_* and applied a stringent Bonferroni-corrected threshold 1.33×10^−4^ (i.e. 0.05/(75×5)) to claim a significant GEI effect.

## URLs

OSCA, http://cnsgenomics.com/software/osca

PLINK2, http://www.cog-genomics.org/plink2

GCTA, http://cnsgenomics.com/software/gcta

UCSC Genome Browser, https://genome.ucsc.edu/

UKB, http://www.ukbiobank.ac.uk/

## Acknowledgements

This research was supported by the Australian Research Council (DP160101343 and DP160101056), the Australian National Health and Medical Research Council (1078037, 1078901, 1113400, 1107258 and 1083656), and the Sylvia & Charles Viertel Charitable Foundation. This study makes use of data from the UK Biobank (project ID: 12514). A full list of acknowledgments of this data set can be found in Supplementary Note 6.

## Author contributions

J.Y. and A.F.M. conceived the study. J.Y., H.W. and A.F.M. designed the experiment. F.Z. developed the software tool. H.W. performed simulations and data analyses under the assistance or guidance from J.Y., J.Z., Y.W., K.K., A.X. and M.Z. J.E.P., M.E.G., N.R.W. and P.M.V. provided critical advice that significantly improved the experimental design and/or interpretation of the results. P.M.V., N.R.W. and J.Y. contributed resources and funding. H.W. and J.Y. wrote the manuscript with the participation of all authors.

