## Supplementary for "Genotype-by-environment interactions inferred from genetic effects on phenotypic variability in the UK Biobank"

Wang *et al.*

**Supplementary Notes 1 to 6**

**Supplementary Figures 1 to 11**

**Supplementary Tables 1 to 8**

**References**

**Supplementary Notes**

### Supplementary Note 1. Bartlett's test, Fligner-Killeen test, and double generalized linear model (DGLM) test

We evaluated four variance quantitative trait locus (vQTL) methods by simulation. Details of the Levene's test have been described in the Methods section of the main text, and details of the other three methods are described below.

The Bartlett's test[1] is one of the earliest methods used to test the inequality of variance but known to be sensitive to the violation of normality assumption[2]. The Bartlett's test-statistic is

$$\frac{(n-k)ln(S_p^2) - \sum_{i=1}^{k}(n_i - 1)ln(S_i^2)}{1 + \frac{1}{3(k-1)}(\sum_{i=1}^{k}(\frac{1}{n_i - 1}) - \frac{1}{n-k})} \sim \chi_{k-1}^2$$

where $n$ is the total sample size; $k$ is the number of groups; $n_i$ is the sample size of the *i*-th group, $n = \sum_{i=1}^{k} n_i$; $S_i^2$ is the sample variance in the *i*-th group; $S_p^2$ is the pooled estimate of the variance, $S_p^2 = \frac{1}{n-k}\sum_{i=1}^{k}(n_i - 1)S_i^2$. We used the *bartlett.test()* function in R.

The Fligner-Killeen (median) test[3] is a rank-based method with similar performance to the Levene's test. The Fligner-Killeen test-statistic is

$$\frac{\sum_{i=1}^{k} n_i\,(\overline{A}_i - \overline{a})^2}{V^2} \sim \chi_{k-1}^2$$

where $n_i$ is the sample size of the *i*-th group, i.e., $n = \sum_{i=1}^{k} n_i$; $\overline{A}_i$ is the mean rank score of the *i*-th group; $\overline{a}$ is the mean rank score of all observations; $V^2$ is the sample variance of rank scores of all observations; "rank score" is assigned by $a = \Phi^{-1}\left(1 + \frac{\frac{i}{n+1}}{2}\right)$ based on the ranks of all observations by $|y_{ij} - \tilde{y}_i|$, and $\tilde{y}_i$ is the median of the *i*-th group. We used the *fligner.test()* function in R.

Ronnegard et al.[4,5] proposed a double generalized linear model (DGLM)[6] that contained two linear models, one for the effect on the trait mean and the other for the effect on the trait variance:

$$E(y|u, u_d) = \mu;\ \mu = Xb + Zu$$

$$var(y|u, u_d) = \phi;\ log(\phi) = X_d b_d + Z_d u_d$$

where *y* is the phenotype; *u* and $u_d$ are the random effects on mean and variance, respectively; *b* and $b_d$ are the fixed effects on mean and variance, respectively. We used "dglm" package in R.

### Supplementary Note 2. Rank-based inverse-normal transformation

We used the simulated data to compare four phenotype processing strategies, i.e., raw phenotype, raw phenotype adjusted for covariates by linear regression, rank-based inverse-normal transformation (RINT) after covariate adjustment, and logarithm transformation after covariate adjustment. RINT was conducted based on the formula below[7,8]

$$y_i^t = \Phi^{-1}\left(\frac{r_i - c}{n - 2c + 1}\right)$$

where $r_i$ is the ordinary rank of the $i$-th observation; $n$ is the total number of observations; $c$ is a constant value (set to 0.5 in this study); $\Phi^{-1}$ is the standard normal quantile function; $y_i^t$ is the transformed value for the $i$-th observation. For RINT after covariate adjustment, we first adjusted the phenotypes for covariates and then transformed the residuals by RINT.

**Supplementary Note 3: The effective number of independent traits**

As some phenotypes were correlated with each other (Supplementary Figure 2), we used an eigendecomposition analysis to estimate the effective number of independent traits[9]. Suppose that **y** is a vector of $p$ phenotypes and **V** is the variance-covariance matrix of vector **y**. The eigen decomposition of matrix **V** is

$$\mathbf{V} = \mathbf{Q}'\mathbf{\Lambda}\mathbf{Q}$$

where **Q** is the matrix of eigenvectors and **Λ** is the diagonal matrix comprised of the ordered eigenvalues $\lambda_1 \dots \lambda_p$. The effective number of $p$ phenotypes can be determined by the formula below[9]:

$$\frac{(\sum_{k=1}^{p} \lambda_k)^2}{\sum_{k=1}^{p} \lambda_k^2}$$

**Supplementary Note 4: Definitions of the three environmental/covariate factors - PA, SB and smoking**

Physical activity (PA) was assessed based on the questions from International Physical Activity Questionnaire (IPAQ)[10], including the number of days per week of walking (DayW), the number of days per week of moderate physical activity (DayW), the number of days per week of vigorous physical activity more than 10 minutes (DayV), the duration of walking (DurW), the duration of moderate physical activity (DurM), and the duration of vigorous physical activity (DurV) (Supplementary Table 3). According to the IPAQ analysis guideline[11], the metabolic equivalents (MET) minutes for walking (METW), moderate physical activity (METM), vigorous physical activity (METV), and the total MET (METT) minutes were calculated by

$$\text{METW} = 3.3 \times \text{DayW} \times \text{DurW}$$

$$\text{METM} = 4.0 \times \text{DayM} \times \text{DurM}$$

$$\text{METV} = 8.0 \times \text{DayV} \times \text{DurV}$$

$$\text{METT} = \text{METW} + \text{METM} + \text{METV}$$

The physical activity level was then labelled as 1) “high” (coded as 3) when “DayV≥3 and METT≥1500” or “DayW+DayM+DayV≥7 and METT≥3000”; 2) “moderate” (coded as 2) when “DayV≥3 and DurV≥20” or “DayM≥5 and DurM≥30” or “DayW≥5 and DurW≥30” or “DayW+DayM+DayV≥5 and METT≥600”; 3) “low” (coded as 1) when no activity or some activity was reported but not enough to meet the criteria above.

Sedentary behaviour (SB) was defined as the sum of the time spent driving (TimeD), non-work-related computer using (TimeC) or TV watching (TimeTV) (Supplementary Table 3). We removed outliers 5 SD from the mean; the remaining data ranged from 0 to 17 hours.

Smoking was assessed based on the answers to two questions about current tobacco smoking (CurS) and past tobacco smoking (PastS) (Supplementary Table 3). Individuals were classified as “never smoker” (coded as 0) if CurS = “no” and PastS = “tried once or twice” or “never”. Individuals were classified as “ever smoker” (coded as 1) if CurS = “most days” or “occasionally”, or PastS = “most days” or “occasionally”.

**Supplementary Note 5: Expected inflation in the Levene's test-statistic due to phantom vQTL effect**

**5.1 Two loci**

Let us consider two genetic loci A and B, and let $p_A$ and $p_B$ denote the frequencies of the major alleles of A and B, respectively, and $p_{AB}$ denote the haplotype frequency of the two major alleles. We know that the LD (including D, D' and $r^2$) between the two loci, the genotype frequencies of the two loci, and the genotype frequency of locus A conditional on locus B are a function of $p_A$, $p_B$ and $p_{AB}$[12].

Allele and haplotype frequencies

| | Locus B | | |
|---|---|---|---|
| Locus A | Major allele B | Minor allele b | Allele frequency |
| Major allele A | $p_{AB}$ | $p_{Ab} = p_A - p_{AB}$ | $p_A$ |
| Minor allele a | $p_{aB} = p_B - p_{AB}$ | $p_{ab} = 1 - p_A - p_B + p_{AB}$ | $p_a = 1 - p_A$ |
| Allele frequency | $p_B$ | $p_b = 1 - p_B$ | 1 |

$p_{AB}$ and LD between A and B as a function of $p_A$ and $p_B$

| Measures | Definition | Maximum value | Minimum value |
|---|---|---|---|
| $p_{AB}$ | - | $min[p_A, p_B]$ | $p_A + p_B - 1$ |
| D | $D = p_{AB} - p_A \times p_B$ | $min[p_A(1-p_B), p_B(1-p_A)]$ | $-(1-p_A)(1-p_B)$ |
| D' | $D' = \frac{D}{\min[p_A(1-p_B), p_B(1-p_A)]}$, if D $> 0$ <br> $D' = \frac{D}{min[p_A p_B, (1-p_A)(1-p_B)]}$, if D $< 0$ | 1 | -1 |
| $r^2$ | $r^2 = \frac{D^2}{p_A p_B (1-p_A)(1-p_B)}$ | $min[\frac{p_A(1-p_B)}{(1-p_A)p_B}, \frac{(1-p_A)p_B}{p_A(1-p_B)}]$ | 0 |

Genotype frequencies of the two loci

| | Genotype BB | Genotype Bb | Genotype bb | Genotype Frequency |
|---|---|---|---|---|
| Genotype AA | $p_{AABB} = p_{AB}^2$ | $p_{AABb} = 2p_{AB}p_{Ab}$ | $p_{AAbb} = p_{Ab}^2 = (p_A - p_{AB})^2$ | $p_A^2$ |

| | | | | |
|---|---|---|---|---|
| | | $= 2p_{AB}(p_A - p_{AB})$ | | |
| Genotype Aa | $p_{AaBB} = 2p_{AB}p_{aB}$<br>$= 2p_{AB}(p_B - p_{AB})$ | $p_{AaBb} = 2(p_{AB}p_{ab} + p_{Ab}p_{aB})$<br>$= 2[p_{AB}(1 - p_A - p_B + p_{AB})$<br>$+(p_A - p_{AB})(p_B - p_{AB})]$ | $p_{Aabb} = 2p_{Ab}p_{ab}$<br>$= 2(p_A - p_{AB})(1 - p_A - p_B$<br>$+ p_{AB})$ | $2p_A(1 - p_A)$ |
| Genotype aa | $p_{aaBB} = p_{aB}^2$<br>$= (p_B - p_{AB})^2$ | $p_{aaBb} = 2p_{aB}p_{ab}$<br>$= 2(p_B - p_{AB})(1 - p_A - p_B$<br>$+ p_{AB})$ | $p_{aabb} = p_{ab}^2$<br>$= (1 - p_A - p_B + p_{AB})^2$ | $(1 - p_A)^2$ |
| Genotype Frequency | $p_B^2$ | $2p_B(1 - p_B)$ | $(1 - p_B)^2$ | 1 |

Genotype frequency of locus A conditioning on locus B

| Genotype | AA | Aa | aa |
|---|---|---|---|
| BB | $P(AA \mid BB) = \frac{p_{AABB}}{p_{BB}}$<br>$= \frac{p_{AB}^2}{p_B^2}$ | $P(Aa \mid BB) = \frac{p_{AaBB}}{p_{BB}}$<br>$= \frac{2p_{AB}(p_B - p_{AB})}{p_B^2}$ | $P(aa \mid BB) = \frac{p_{aaBB}}{p_{BB}}$<br>$= \frac{(p_B - p_{AB})^2}{p_B^2}$ |
| Bb | $P(AA \mid Bb) = \frac{p_{AABb}}{p_{Bb}}$<br>$= \frac{p_{AB}(p_A - p_{AB})}{p_B(1 - p_B)}$ | $P(Aa \mid Bb) = \frac{p_{AaBb}}{p_{Bb}}$<br>$= \frac{p_{AB}(1 - p_A - p_B + p_{AB}) + (p_A - p_{AB})(p_B - p_{AB})}{p_B(1 - p_B)}$ | $P(aa \mid Bb) = \frac{p_{aaBb}}{p_{Bb}}$<br>$= \frac{(p_B - p_{AB})(1 - p_A - p_B + p_{AB})}{p_B(1 - p_B)}$ |
| bb | $P(AA \mid bb) = \frac{p_{AAbb}}{p_{bb}}$<br>$= \frac{(p_A - p_{AB})^2}{(1 - p_B)^2}$ | $P(Aa \mid bb) = \frac{p_{Aabb}}{p_{bb}}$<br>$= \frac{2(p_A - p_{AB})(1 - p_A - p_B + p_{AB})}{(1 - p_B)^2}$ | $P(aa \mid bb) = \frac{p_{aabb}}{p_{bb}}$<br>$= \frac{(1 - p_A - p_B + p_{AB})^2}{(1 - p_B)^2}$ |

#### 5.2 Causal variant (locus A) with an additive genetic effect

Let us assume that locus A is the causal variant with an additive genetic effect ($b_c$) on a phenotype:

$$y \sim (\mu + b_c x_a, \sigma^2)$$

| Genotype | Code ($x_a$) | $E(y/x_a)$ | $Var(y/x_a)$ | $E(y^2 \mid x_a)$ |
|---|---|---|---|---|
| AA | 0 | $\mu$ | $\sigma^2$ | $\sigma^2 + \mu^2$ |
| Aa | 1 | $\mu + b_c$ | $\sigma^2$ | $\sigma^2 + (\mu + b_c)^2$ |
| aa | 2 | $\mu + 2b_c$ | $\sigma^2$ | $\sigma^2 + (\mu + 2b_c)^2$ |

The expected phenotypic mean and variance given a genotype of locus B (marker) can be found in the tables below.

| Genotype | Code ($x_b$) | *E(y\|x_b)* |
|---|---|---|
| BB | 0 | $\mu + \frac{2b_c(p_B - p_{AB})}{p_B}$ |
| Bb | 1 | $\mu + \frac{b_c(2p_B - p_{AB} - 2p_B^2 + 2p_B p_{AB} - p_A p_B)}{p_B(1-p_B)}$ $= \mu + \frac{b_c[(p_B - p_{AB})(1-p_B) + (1 - p_A - p_B + p_{AB})p_B]}{p_B(1-p_B)}$ |
| bb | 2 | $\mu + \frac{2b_c(1 - p_A - p_B + p_{AB})}{1 - p_B}$ |

| *Var(y\|x_b)* |
|---|
| $\sigma^2 + \frac{2{b_c}^2(p_B - p_{AB})p_{AB}}{p_B^2}$ |
| $\sigma^2 + \frac{{b_c}^2(p_B p_{AB} - p_{AB}^2 + 2p_B p_{AB}^2 - 3p_B^2 p_{AB} + p_A p_B^2 - 2p_B^2 p_{AB}^2 + 2p_A p_B^2 p_{AB} + 2p_B^3 p_{AB} - p_A p_B^3 - p_A^2 p_B^2)}{p_B^2(1-p_B)^2}$ $= \sigma^2 + \frac{{b_c}^2[(p_B - p_{AB})p_{AB}(1-p_B)^2 + (1 - p_A - p_B + p_{AB})(p_A - p_{AB})p_B^2]}{p_B^2(1-p_B)^2}$ |
| $\sigma^2 + \frac{2{b_c}^2(1 - p_A - p_B + p_{AB})(p_A - p_{AB})}{(1-p_B)^2}$ |

We therefore can observe an additive effect on both mean ($b_m$) and variance ($\beta_m$) at the marker (locus B):

$$y \sim (\mu + b_m x_b, \sigma^2 + \beta_m x_b)$$

where

- $b_m = \frac{b_c(p_{AB} - p_A p_B)}{p_B(1 - P_B)}$
- $\beta_m = \frac{b_c^2[(1-2p_B)p_{AB}^2 + (2p_A p_B + p_B - 1)p_B p_{AB} + (1 - p_A - p_B)p_A p_B^2]}{p_B^2(1-p_B)^2}$

**5.3 QTL test-statistics at the marker variant (locus B)**

Assuming phenotypic variance of 1 (i.e., var($y$) = 1), the variance explained by the marker variant ($q_m^2$) and the non-centrality parameter (NCP) of a chi-squared test for QTL effect at the marker can be written as

- $q_m^2 = 2p_B(1-p_B)b_m^2 = 2p_B(1-p_B)\frac{b_c^2(p_{AB}-p_Ap_B)^2}{p_B^2(1-p_B)^2} = 2p_A(1-p_A)b_c^2\frac{(p_{AB}-p_Ap_B)^2}{p_A(1-p_A)p_B(1-p_B)} = q_c^2r^2$
- $\text{NCP} = \frac{nq_m^2}{1-q_m^2} = \frac{nq_c^2r^2}{1-q_c^2r^2}$

where $n$ is the sample size, $q_c^2$ is the variance explained by the causal variant, and $r^2$ is the LD between the causal and the marker variants. This derivation is consistent with that in previous studies[13,14].

#### 5.4 vQTL test statistic at the marker variant (locus B)

Under normality assumption, the distribution of the phenotype with respect to the marker variant can be written as:

$$y \sim N(\mu + b_m x_b, \sigma^2 + \beta_m x_b)$$

We then have

$$y - E(y|x_b) \sim N(0, \sigma^2 + \beta_m x_b)$$

, and $z = |y - \widetilde{y}\,|$

$$z = |y - \widetilde{y}\,| = |y - E(y|x_b)|$$
$$\sim \text{ Folded Normal Distribution}(\sqrt{\frac{2}{\pi}(\sigma^2 + \beta_m x_b)}, (1-\frac{2}{\pi})(\sigma^2 + \beta_m x_b))$$

| Genotype | Code ($x_b$) | E($z$\|$x_b$) | var($z$\|$x_b$) | $E(z^2\|x_b)$ |
|---|---|---|---|---|
| BB | 0 | $\sqrt{\frac{2}{\pi}\sigma^2}$ | $(1-\frac{2}{\pi})\sigma^2$ | $\sigma^2$ |
| Bb | 1 | $\sqrt{\frac{2}{\pi}(\sigma^2+\beta_m)}$ | $(1-\frac{2}{\pi})(\sigma^2+\beta_m)$ | $\sigma^2+\beta_m$ |

| | | | | |
|---|---|---|---|---|
| bb | 2 | $\sqrt{\frac{2}{\pi}(\sigma^2 + 2\beta_m)}$ | $(1 - \frac{2}{\pi})(\sigma^2 + 2\beta_m)$ | $\sigma^2 + 2\beta_m$ |

$$E(z) = E(z|x_b = 0)P(x_b = 0) + E(z|x_b = 1)P(x_b = 1) + E(z|x_b = 2)P(x_b = 2)$$
$$= \sqrt{\frac{2}{\pi}\sigma^2 {p_B}^2} + \sqrt{\frac{2}{\pi}(\sigma^2 + \beta_m)2p_B(1 - p_B)} + \sqrt{\frac{2}{\pi}(\sigma^2 + 2\beta_m)(1 - p_B)^2}$$

$$E(z^2) = E(z^2|x_b = 0)P(x_b = 0) + E(z^2|x_b = 1)P(x_b = 1) + E(z^2|x_b = 2)P(x_b = 2)$$
$$= \sigma^2 {p_B}^2 + (\sigma^2 + \beta_m)2p_B(1 - p_B) + (\sigma^2 + 2\beta_m)(1 - p_B)^2$$
$$= \sigma^2 + 2(1 - p_B)\beta_m$$

$$var(z) = E(z^2) - [E(z)]^2 = \sigma^2 + 2(1 - p_B)\beta_m - [E(z)]^2$$

The Levene's test is essentially one-way ANOVA test on the variable $z$ (see the Methods section). We therefore have

$$E(SST) = E[\sum_{i=1}^{k}\sum_{j=1}^{n_i}(z_{ij} - z_{..})^2] = Var(z)n = (\sigma^2 + 2(1 - p_B)\beta_m - [E(z)]^2)n;$$

$$E(SSE) = E\left[\sum_{i=1}^{k}\sum_{j=1}^{n_i}(z_{ij} - z_{i.})^2\right]$$
$$= (1 - \frac{2}{\pi})\sigma^2 n{p_B}^2 + (1 - \frac{2}{\pi})(\sigma^2 + \beta_m)n2p_B(1 - p_B) + (1 - \frac{2}{\pi})(\sigma^2 + 2\beta_m)(1 - p_B)^2$$
$$= (1 - \tfrac{2}{\pi})(\sigma^2 + 2(1 - p_B)\beta_m)n;$$

$$E(SSR) = E(SST - SSE) = [\tfrac{2}{\pi}(\sigma^2 + 2(1 - p_B)\beta_m) - [E(z)]^2]n;$$

$$F_{\text{Levene}} = \frac{(n-3)E(SSR)}{(3-1)E(SSE)} \approx \frac{n}{2}\frac{E(SSR)}{E(SSE)} = \frac{n}{2}\frac{\frac{2}{\pi}(\sigma^2 + 2(1 - p_B)\beta_m) - [E(z)]^2}{(1 - \frac{2}{\pi})(\sigma^2 + 2(1 - p_B)\beta_m)}$$
$$= \frac{n}{\pi - 2}(1 - \frac{[\sqrt{\sigma^2}{p_B}^2 + \sqrt{\sigma^2 + \beta_m}2p_B(1 - p_B) + \sqrt{\sigma^2 + 2\beta_m}(1 - p_B)^2]^2}{\sigma^2 + 2(1 - p_B)\beta_m})$$

where $F_{\text{Levene}}$ is the Levene's $F$-statistic; $SST$, $SSR$ and $SSR$ are the total sum of squares, regression sum of squares and error sum of squares, respectively, as defined in an ANOVA analysis.

Given that var(y) = 1, we can replace $b_c^2$ with $\frac{q_c^2}{2p_A(1-p_A)}$, and $\sigma^2$ with $1 - q_c^2$:

$$\beta_m = \frac{q_c^2[(1-2p_B)p_{AB}^2 + (2p_Ap_B + p_B - 1)p_Bp_{AB} + (1 - p_A - p_B)p_Ap_B^2]}{2p_A(1-p_A)p_B^2(1-p_B)^2}$$

$$F_{\text{Levene}} = \frac{n}{\pi - 2}(1 - \frac{[\sqrt{1-q_c^2}{p_B}^2 + \sqrt{1-q_c^2+\beta_m}2p_B(1-p_B) + \sqrt{1-q_c^2+2\beta_m}(1-p_B)^2]^2}{1-q_c^2+2(1-p_B)\beta_m})$$

Therefore, the phantom vQTL test statistic is a function of sample size $n$, variance explained by the causal variant $q_c^2$, allele frequency of the causal variant $p_A$, allele frequency of the marker variant $p_B$, and the haplotype frequency $p_{AB}$. This formula has been confirmed by simulation (Supplementary Figure 7).

#### 5.5 The maximum test statistics of phantom vQTLs given a pair of causal and marker variants

Since $|\beta_m|$ increases monotonically with the increases of $F_{\text{Levene}}$, finding the maximum value of $F_{\text{Levene}}$ is equivalent to finding the maximum value of $|\beta_m|$. We know from the derivation above that $\beta_m$ is a quadratic function of $p_{AB}$ so that $|\beta_m|$ is maximised (considering that $\beta_m$ can be negative) when $p_{AB}$ is

- the minimum value $p_A + p_B - 1$ (i.e., LD D'=-1);
- or the stationary point $\frac{p_B(2p_Ap_B+p_B-1)}{2(2p_B-1)}$ if available;
- or the maximum value $min[p_A, p_B]$ (i.e., LD D'=1).

We computed $F_{\text{Levene}}$ given a number of parameters including $p_{AB}$ (the three possible maximum values mentioned above), $p_a$ (ranging from 0.001 to 0.5, equivalent to $p_A$ from 0.999 to 0.5), $p_b$ (ranging from 0.05 to 0.5, equivalent to $p_B$ from 0.95 to 0.5), $q_c^2$ (= 0.005, 0.01 or 0.02) and $n$ (= 350,000) (Supplementary Figure 8).

**Supplementary Note 6. Acknowledgements**

This study has been conducted using the UK Biobank resource under Application Number 12505. The UK Biobank was established by the Wellcome Trust medical charity, Medical Research Council, Department of Health, Scottish Government and the Northwest Regional Development Agency. It has also had funding from the Welsh Assembly Government, British Heart Foundation and Diabetes UK.

### Supplementary Figures

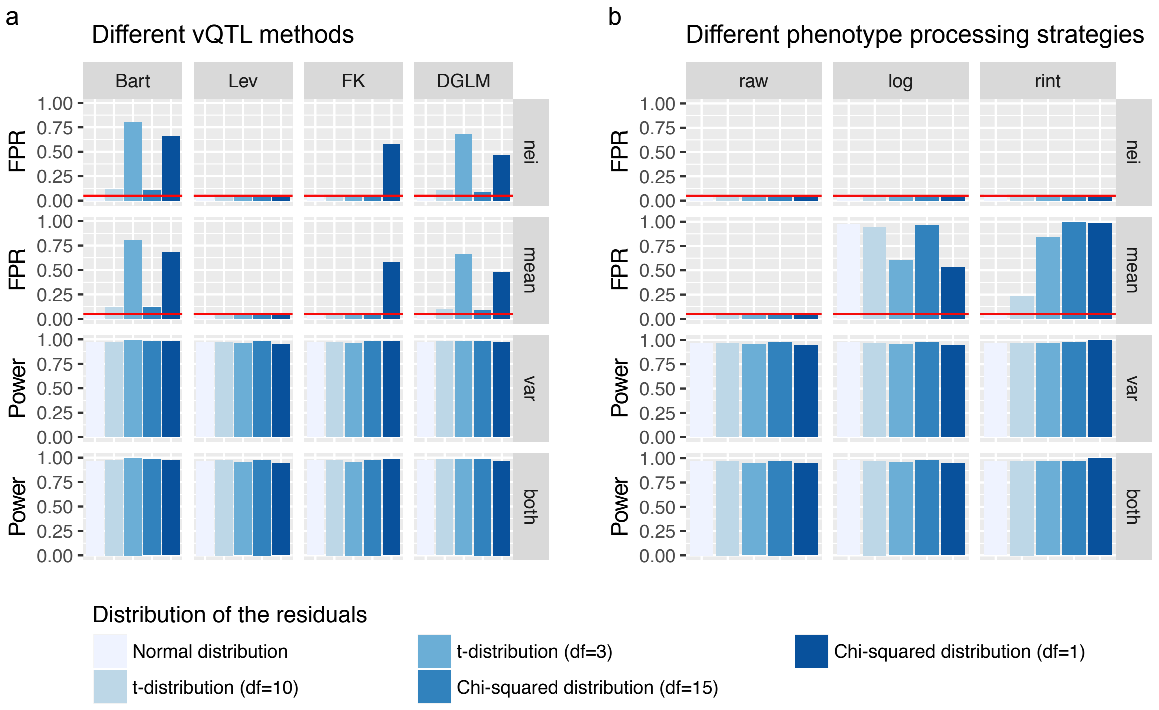


**Supplementary Figure 1. Evaluation of statistical methods (a) and phenotype processing strategies (b) for vQTL analysis by simulation based on a single-SNP model.** Phenotypes of 10,000 individuals were simulated based on one SNP and one error term in a single-SNP model (Methods). The SNPs effects were simulated under four scenarios: 1) effect on neither mean nor variance (nei), 2) effect on mean only (mean), 3) effect on variance only (var), or 4) effect on both mean and variance (both). The error term was generated from 5 different distributions: normal distribution, $t$-distribution with degree of freedom (df) = 10 or 3, or $\chi^2$ distribution with df = 15 or 1. Four statistical test methods, i.e. the Bartlett's test (Bart), the Levene's test (Lev), the Fligner-Killen test (FK) and the DGLM, were used to detect vQTLs. In panel b, the Levene's test was used to analyse phenotypes processed using three strategies, i.e. raw phenotype (raw), rank-based inverse-normal transformation (rint), and logarithm transformation (log). The FPR or power was calculated as the number of vQTLs with $p < 0.05$ divided by the total number of tests across 1,000 simulations. The red horizontal line represents an FPR of 0.05.

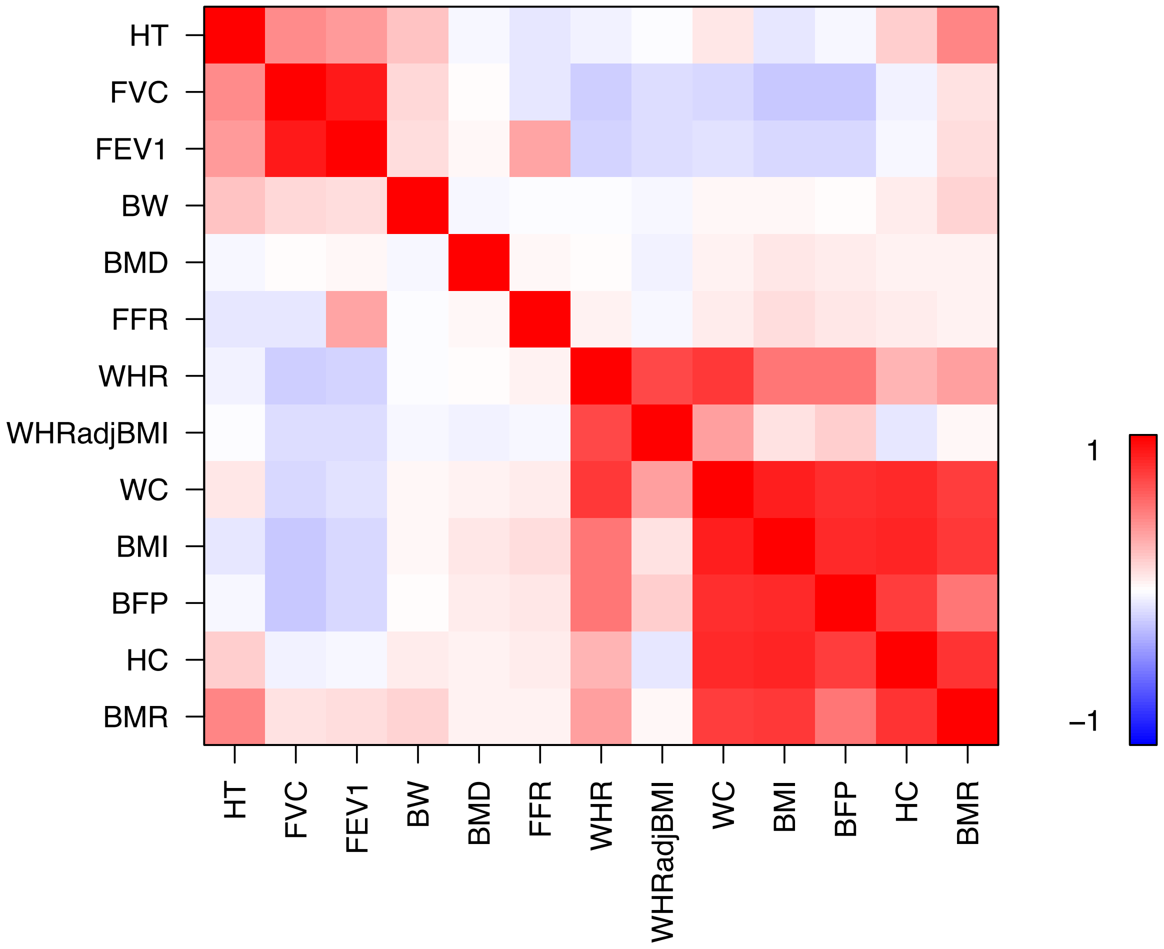


**Supplementary Figure 2. Phenotypic correlations among the 13 UKB traits.** The Pearson's correlation coefficient was calculated between each pair of the processed phenotypes. The order of the traits shown on the plot above was determined by hierarchical cluster analysis using the R function *hclust()*.

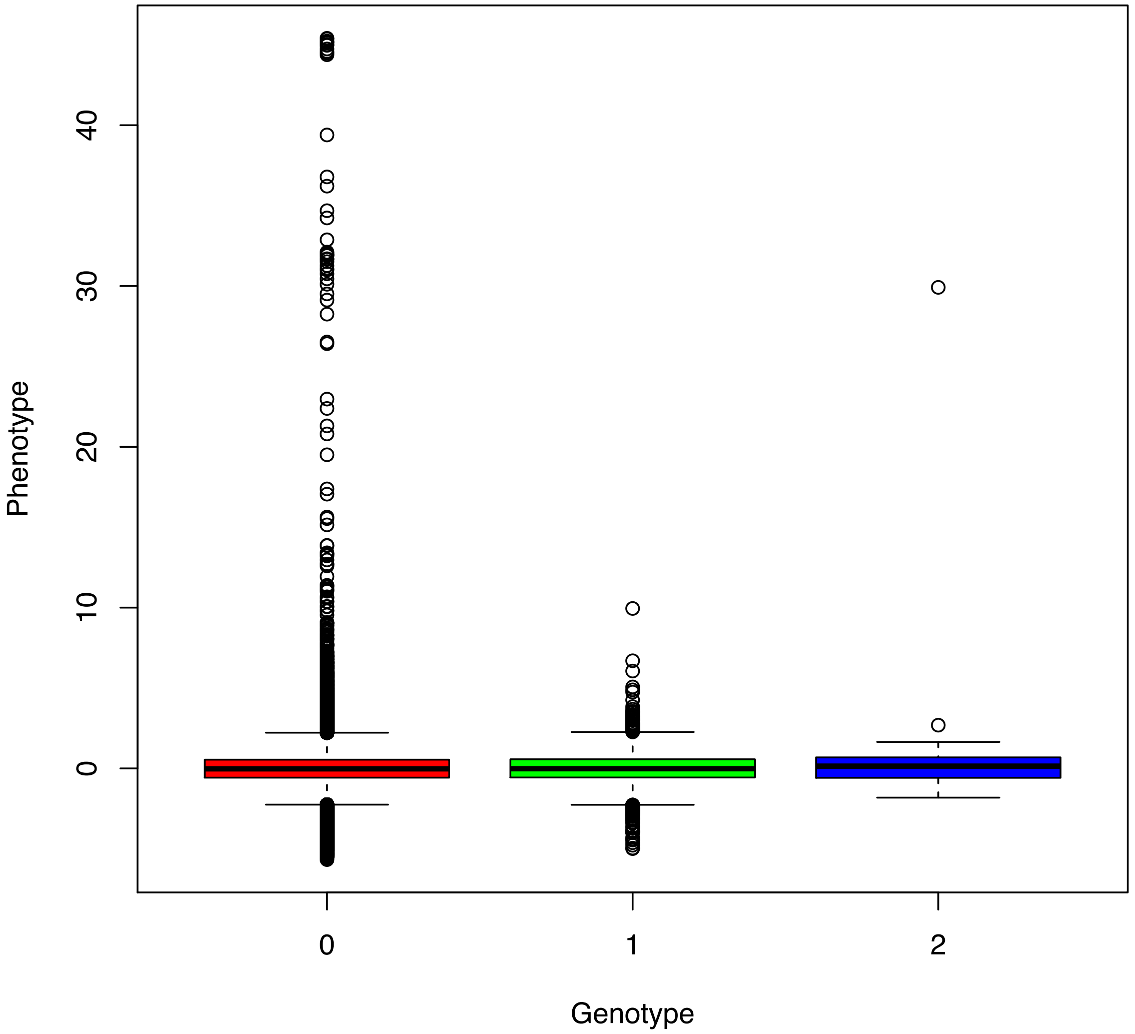


**Supplementary Figure 3. Spurious vQTL association due to the coincidence of a minor allele with a phenotypic outlier.** This is an example that a spurious vQTL signal ($P_{\mathrm{vQTL}}$ = $4.48\times10^{-9}$) at a low-MAF variant (MAF = 0.012) is caused by the coincidence of a minor allele with a phenotypic outlier for FVC. The variance of the phenotype (after covariates adjustment and standardisation) are 1.00, 0.83 and **20.20** in the three genotype groups of rs11102024 respectively. Note that for all the other vQTL results presented in this paper are from analyses excluding individuals with adjusted phenotypes more than 5 SD from the mean and SNPs with MAF < 0.05.

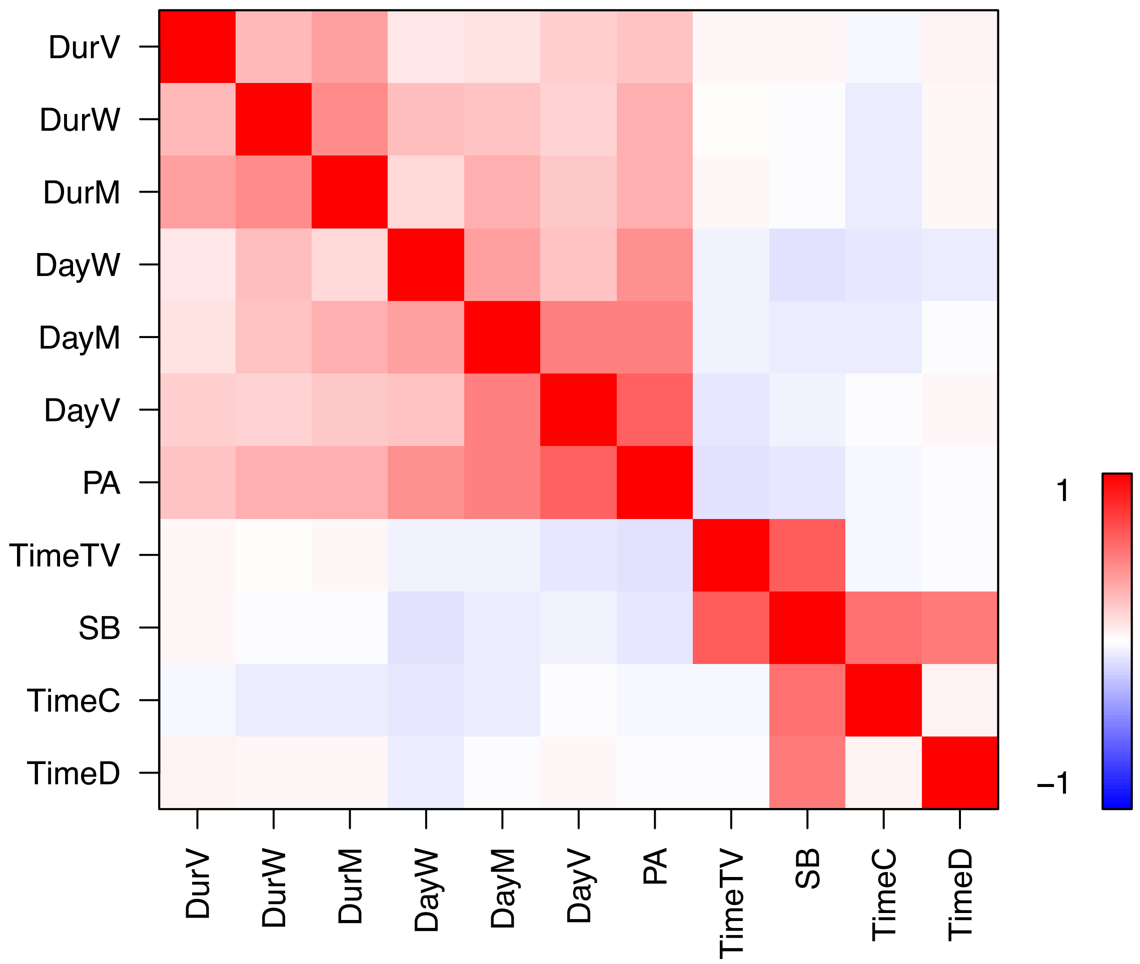


**Supplementary Figure 4. Phenotypic correlations among PA and SB measures in the UKB.** The Pearson's correlation coefficient was calculated between each pair of the measures. The order of the traits shown on the plot above was determined by hierarchical cluster analysis using the R function *hclust()*.

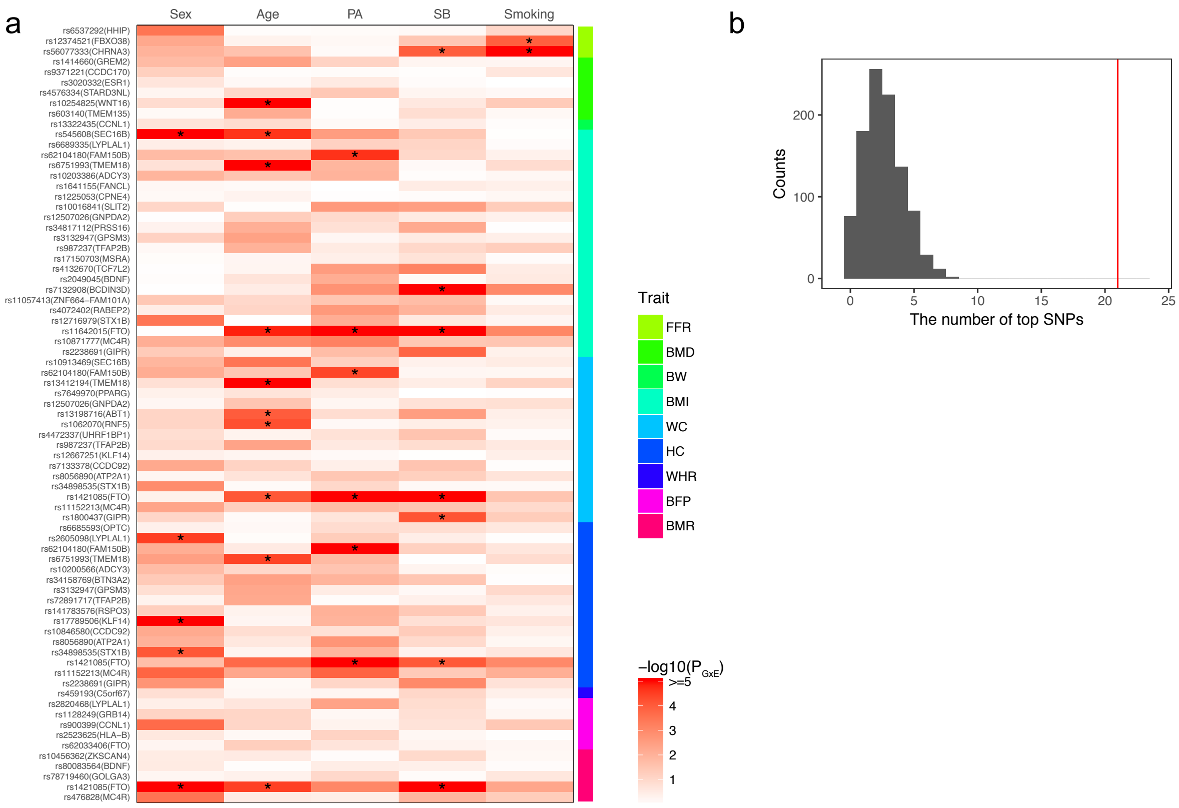


**Supplementary Figure 5. Enrichment of GEI effects among the 75 vQTLs in compared with a random set of QTLs using the raw phenotypic values.** Five environmental factors, i.e., sex, age, physical activity (PA), sedentary behaviour (SB), and smoking, were used in the GEI analysis. (a) The heatmap plot of GEI test statistics (-$\log_{10}(P_{\mathrm{GEI}})$) for the 75 top vQTL SNPs. "*" denotes significant GEI effects after Bonferroni correction ($P_{\mathrm{GEI}} < 1.33\times10^{-4} = 0.05/75/5$). (b) The distribution of the number of significant GEI effects for 75 top QTL SNPs randomly selected from all the top QTL SNPs with 1000 repeats (mean 2.60 and SD 1.57). The red line represents the number of significant GEI effects for the 75 top vQTL SNPs (i.e., 21).

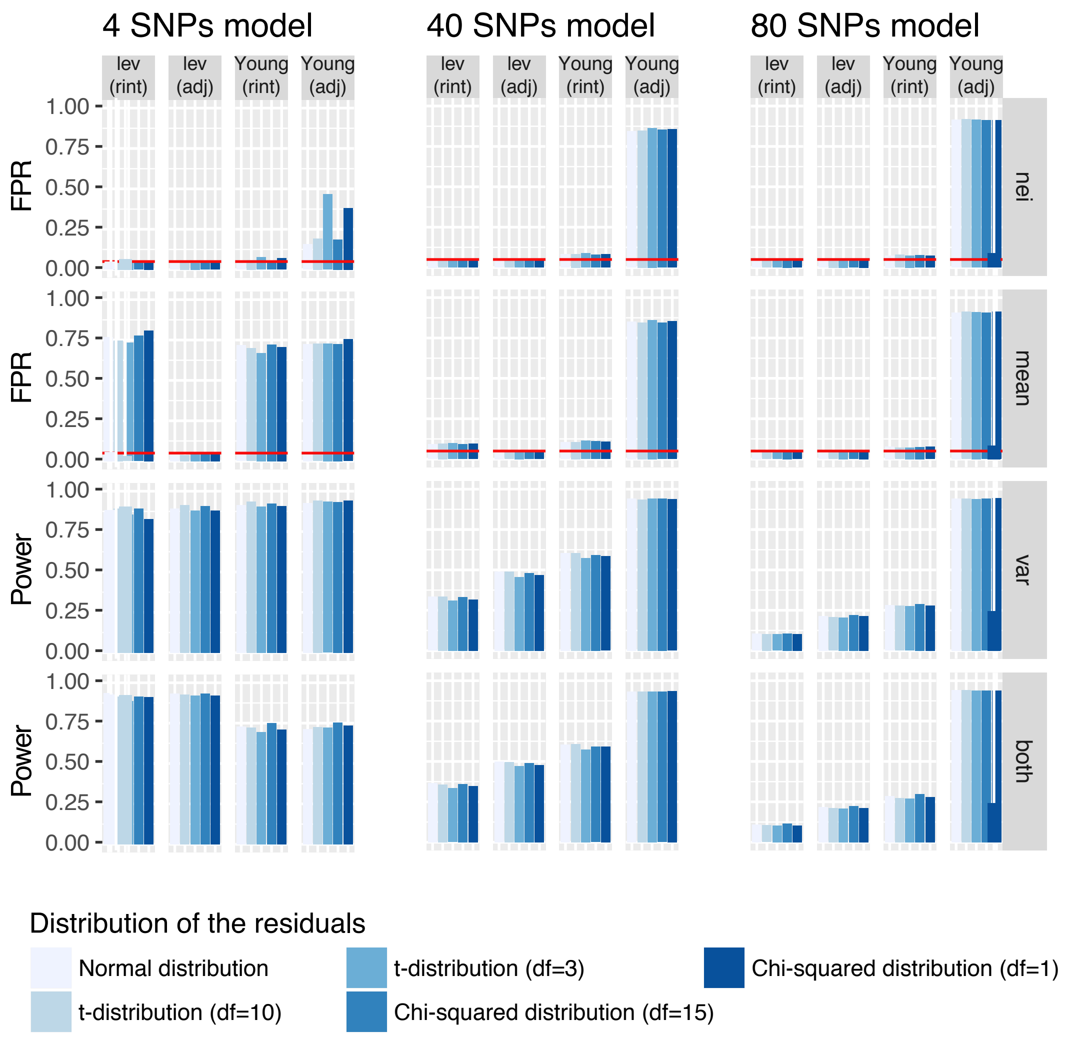


**Supplementary Figure 6. Comparison of the Young et al. method with the Levene's test by vQTL simulation.** While we were preparing the manuscript, a very recent study from Young et al.[15] developed an efficient algorithm for fitting DGLM (called heteroskedastic linear mixed model or HLMM) and proposed a dispersion effect test (DET) to remove the impact of the QTL effects on the vQTL signals. We used our multiple-SNP simulation setting (Figure 1 and Methods) to quantify the FPR and power of the Young et al. method (HLMM + DET) in comparison with the Levene's test based on the phenotype after 1) covariate adjustment ("adj") or 2) covariate adjustment followed by rank-based inverse-normal transformation ("rint"). For the Levene's test, the FPR or power was computed as the number of vQTLs with $p < 0.05$ divided by the total number of tests across 1,000 simulations. For the analysis with the Young et al. method, the FPR or power was computed as the number of vQTLs with DET $p < 0.05$ divided by the total number of tests across simulations.

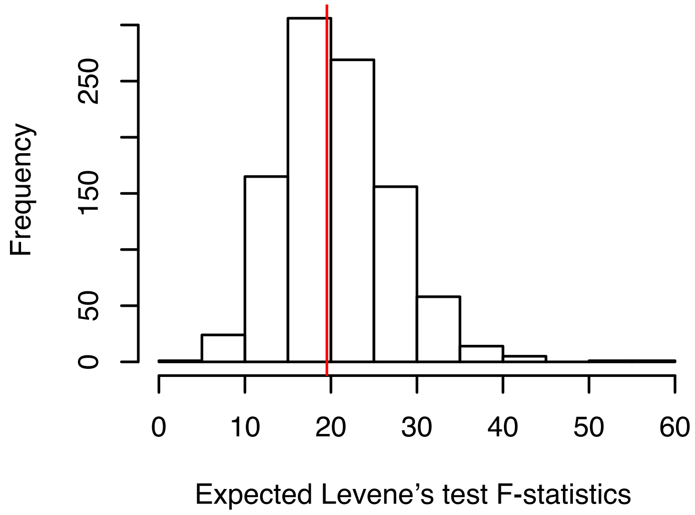


**Supplementary Figure 7. Verification of the expected Levene's test *F*-statistic due to phantom vQTL effect by simulation.** We simulated two variants A ($p_A$ = 0.7) and B ($P_B$ = 0.6) in LD ($P_{AB}$ = 0.6, LD $r^2$ = 0.64, and LD D' = 1) from multinomial(2, ($P_{AB}$, $P_{Ab}$, *PaB*, *Pab*)) and a phenotype based on the causal variant A explaining 5% variance in 350,000 individuals. Shown is the distribution of *F*-statistics from the Levene's test using the simulated data with 1,000 replicates. The red line indicates the theoretical value based on the formula in Supplementary Note 5.4.

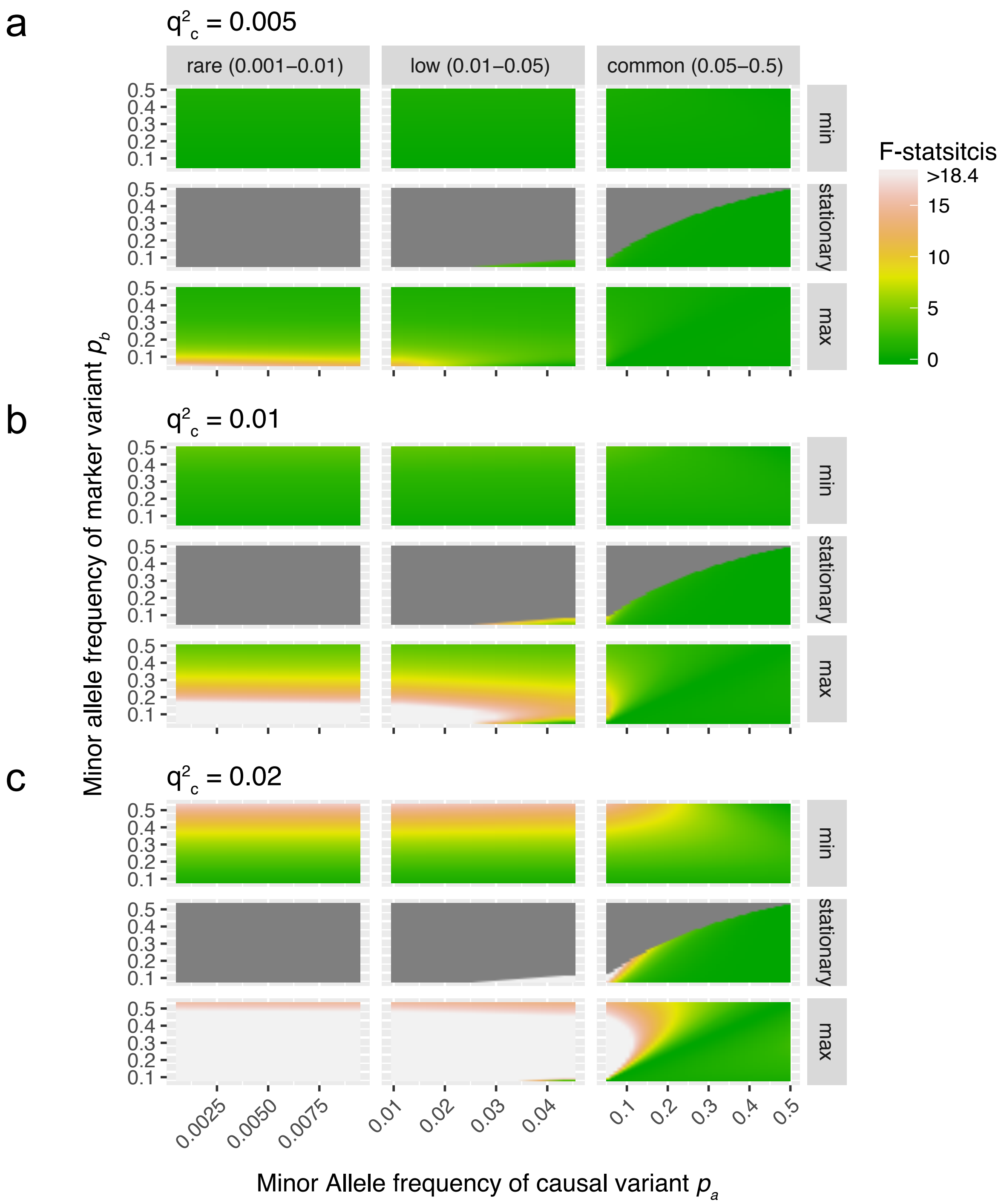


**Supplementary Figure 8. Expected phantom vQTL *F*-statistics from the Levene's test.** We calculated the expected phantom vQTL *F*-statistics given a number of parameters including $p_{AB}$ (three possible maximum values), $p_a$ (ranging from 0.001 to 0.5), $p_b$ (ranging from 0.05 to 0.5), $q_c^2$ (= 0.005, 0.01 or 0.02) and $n$ (= 350,000) (Supplementary Note 5 for more details). An $F$ value of 18.4 is equivalent to a genome-wide significant p-value of $1{\times}10^{-8}$.

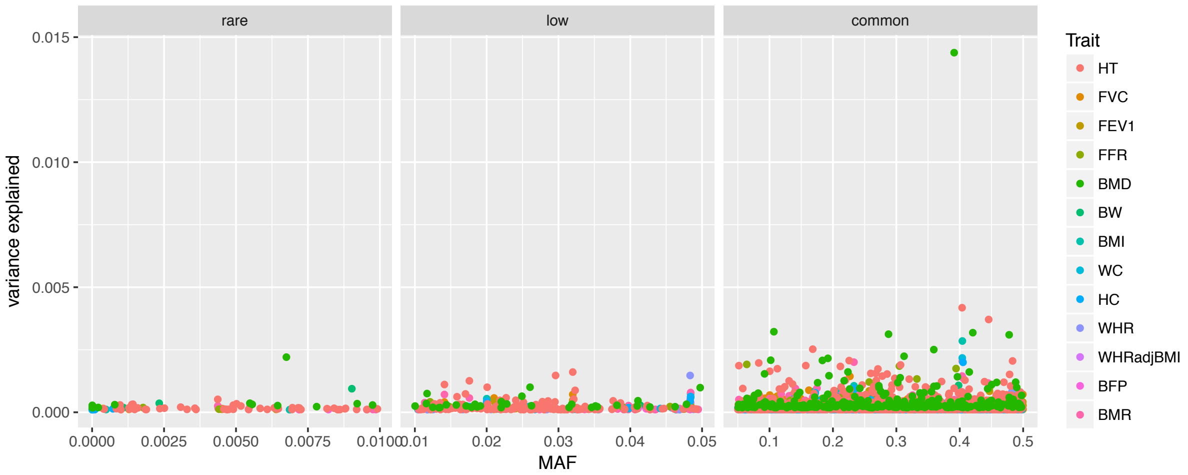


**Supplementary Figure 9. Estimated variance explained by top QTL SNPs for the 13 UKB traits.** Note that because the phantom vQTL signals at common SNPs can be induced by rare (MAF≤0.01) or low-frequency (0.01≤MAF<0.05) variants, we extended our GWAS analysis to all 44,741,800 imputed variants (MAF<0.05). The estimated variance explained by each GWAS top SNP is plotted against its MAF.

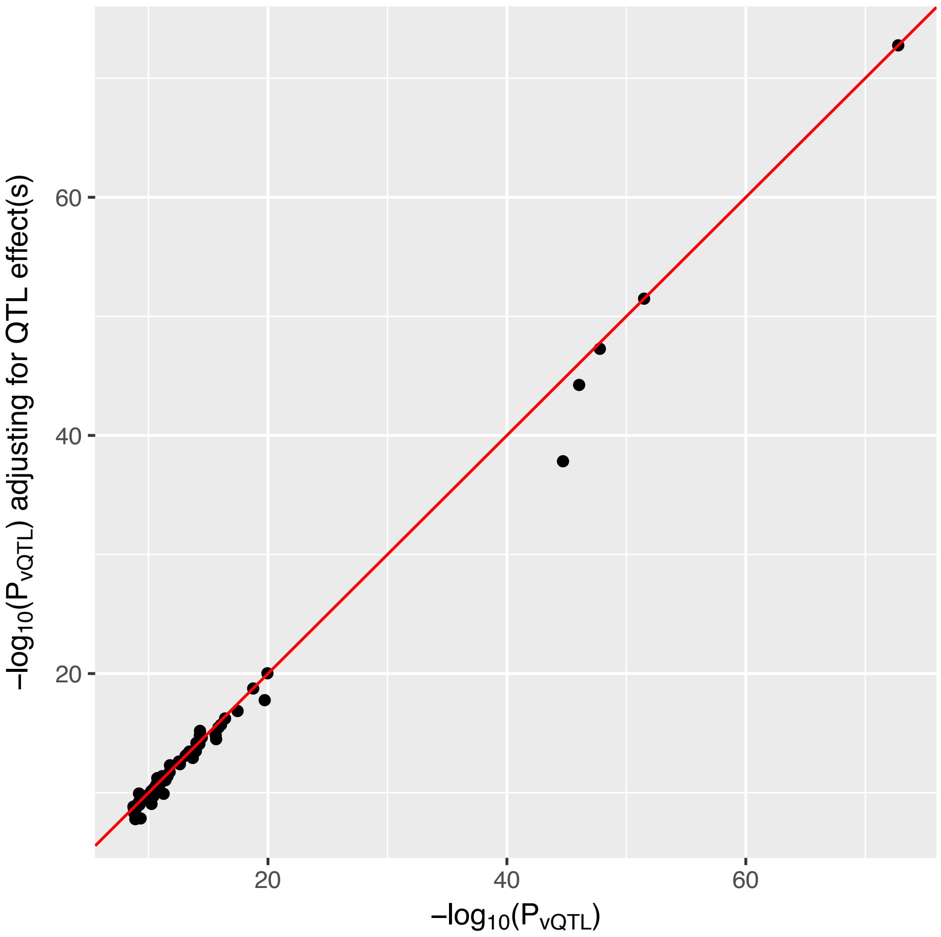


**Supplementary Figure 10. vQTL test statistics ($-\log_{10}(P_{vQTL})$) from analyses with and without adjusting the phenotype for the QTL effect(s) of the top GWAS SNP(s) within 10Mb of the top vQTL SNP.** The red line represents the line with slope 1 and intercept 0.

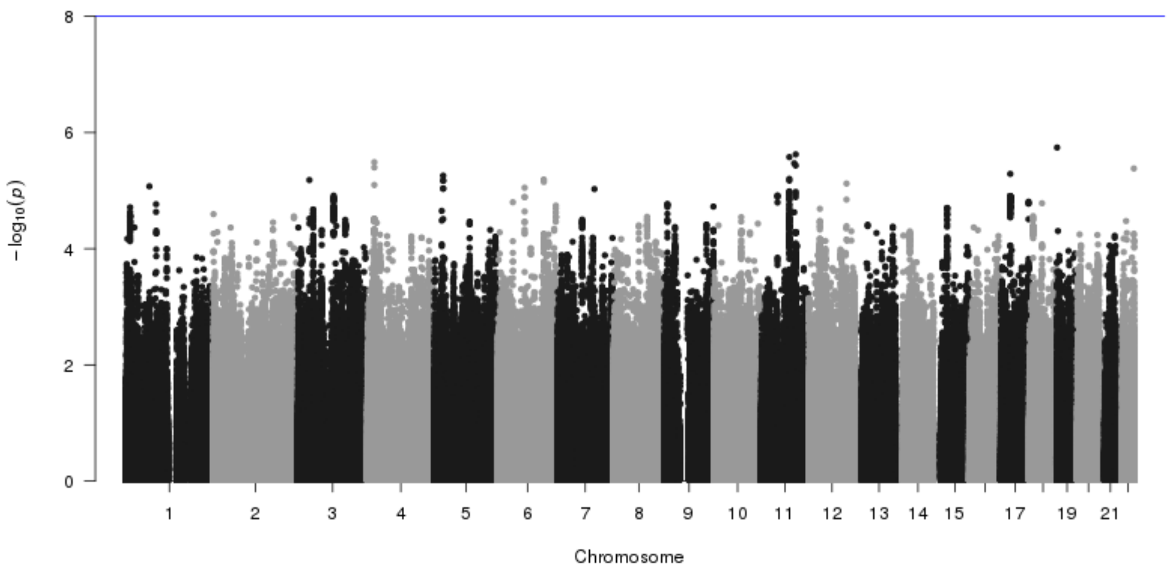


**Supplementary Figure 11. Manhattan plot of epistasis analysis for one of top vQTL SNPs.** We conducted epistasis analysis between each of 75 top vQTL SNPs and any other SNPs in more than 10 Mb distance or on a different chromosome for the relevant trait using PLINK2[16] (--epistasis option). The blue horizontal line represents the genome-wide significance level (i.e., p-value = $1 \times 10^{-8}$). Shown are the results from the epistasis analysis with the top vQTL SNP rs10913469 for waist circumference (WC).

### Supplementary Tables

#### Supplementary Table 1. Descriptive summary of the 13 UKB traits

| Trait | Description | Sample size | Distribution of raw phenotype | Distribution of processed phenotype | UDI[a] |
|---|---|---|---|---|---|
| HT | Standing height | 347,086 | 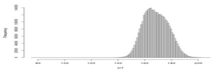 | 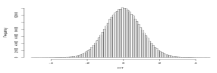 | 50-0.0 |
| FVC | Forced vital capacity | 317,222 | 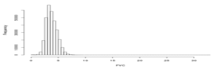 | 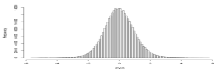 | 3062-0.0 |
| FEV1 | Forced expiratory volume in 1-second | 317,285 | 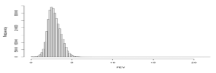 | 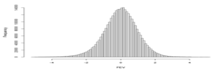 | 3063-0.0 |
| FFR[b] | FEV1 and FVC ratio | 316,614 | 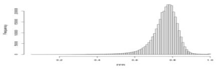 | 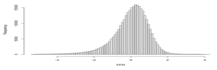 | NA |
| BMD | Heel bone mineral density T-score, automated | 197,261 | 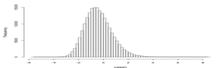 | 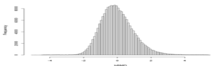 | 78-0.0 |
| BW | Birth weight | 197,758 | 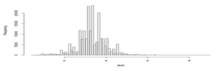 | 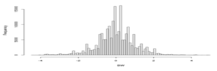 | 20022-0.0 |
| BMI | Body mass index (BMI) | 346,393 | 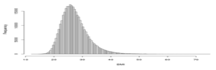 | 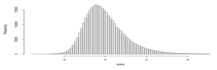 | 21001-0.0 |
| WC | Waist circumference | 347,158 | 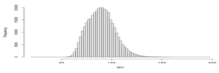 | 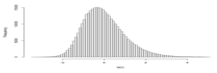 | 48-0.0 |
| HC | Hip circumference | 346,781 | 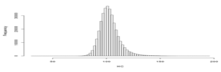 | 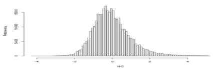 | 49-0.0 |
| WHR[c] | Waist to Hip Ratio | 347,134 | 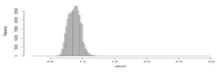 | 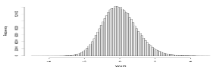 | NA |
| WHRadjBMI[d] | WHR adjusted for BMI | 177,060 | 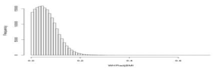 | 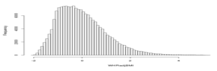 | NA |
| BFP | Body fat percentage | 341,632 | 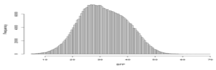 | 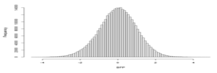 | 23099-0.0 |
| BMR | Basal metabolic rate | 341,584 | 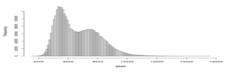 | 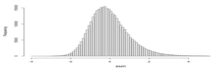 | 23105-0.0 |

Note: a) UDI, the Unique Data Identifier in the UKB dataset; b) FFR is the ratio of FEV1 to FVC; c) WHR is the ratio of waist circumference to hip circumference; d) WHRadjBMI is the residual after adjusting WHR for BMI.

**Supplementary Table 2. Seventy-five experiment-wise significant vQTLs for 9 UKB traits.**

| Trait | CHR | SNP | bp | Nearest Gene | MAF | vQTL p-value | QTL p-value | Phenotypic variance in each genotype group | Phenotypic mean in each genotype group |
|---|---|---|---|---|---|---|---|---|---|
| FFR | 4 | rs6537292 | 145469968 | HHIP | 0.394 | 1.97E-14 | 3.58E-122[a] | 1.0217,0.9936,0.9561 | -0.0453,0.0091,0.0787 |
| | 5 | rs12374521 | 147836880 | FBXO38 | 0.456 | 7.10E-10 | 1.60E-58 | 1.0223,0.9978,0.97 | -0.039,0.0055,0.0417 |
| | 15 | rs56077333 | 78899003 | CHRNA3 | 0.325 | 1.09E-14 | 2.11E-06 | 0.9757,1.0107,1.0588 | 0.0072,-0.0019,-0.0225 |
| BMD | 1 | rs1414660 | 240586695 | GREM2 | 0.192 | 7.83E-14 | 1.28E-94 | 0.977,1.0362,1.0452 | -0.0322,0.0523,0.1304 |
| | 6 | rs9371221 | 151885986 | CCDC170 | 0.101 | 4.59E-10 | 1.30E-76 | 1.0097,0.9502,0.9408 | 0.02,-0.0817,-0.1479 |
| | 6 | rs3020332 | 152008924 | ESR1 | 0.45 | 5.42E-14 | 8.94E-130 | 0.966,0.997,1.0429 | -0.074,0.0126,0.0795 |
| | 7 | rs4576334 | 38153747 | STARD3NL | 0.196 | 2.36E-13 | 2.60E-86 | 0.9784,1.0308,1.0684 | -0.0325,0.0511,0.1152 |
| | 7 | rs10254825 | 120956440 | WNT16 | 0.391 | 2.01E-45 | 0 | 0.9279,1.0107,1.057 | -0.1455,0.05,0.1903 |
| | 11 | rs603140 | 86884615 | TMEM135 | 0.312 | 1.61E-12 | 4.48E-98 | 1.0149,0.9924,0.9333 | 0.0417,-0.0204,-0.1142 |
| BW | 3 | rs13322435 | 156795468 | CCNL1 | 0.402 | 9.71E-10 | 6.21E-48 | 1.0287,0.9847,0.9742 | 0.0376,-0.0072,-0.0585 |
| BMI | 1 | rs545608 | 177899121 | SEC16B | 0.206 | 3.88E-17 | 1.97E-63 | 0.9801,1.0251,1.0835 | -0.0202,0.0282,0.0847 |
| | 1 | rs6689335 | 219628682 | LYPLAL1 | 0.419 | 2.86E-12 | 4.73E-08 | 1.0249,0.9907,0.972 | 0.0106,-0.0013,-0.0167 |
| | 2 | rs62104180 | 466003 | FAM150B | 0.05 | 1.22E-11 | 3.57E-51 | 1.0054,0.9461,0.8598 | 0.0083,-0.075,-0.1488 |
| | 2 | rs6751993 | 635864 | TMEM18 | 0.167 | 3.50E-18 | 3.31E-65 | 1.0155,0.9707,0.9188 | 0.0197,-0.0361,-0.0912 |
| | 2 | rs10203386 | 25136866 | ADCY3 | 0.452 | 1.33E-11 | 8.45E-43 | 0.9768,0.9994,1.0333 | -0.0272,3e-04,0.0404 |
| | 2 | rs1641155 | 58965211 | FANCL | 0.311 | 1.25E-09 | 4.42E-17 | 0.9872,1.0092,1.0266 | -0.0141,0.0108,0.0266 |
| | 3 | rs1225053 | 131642852 | CPNE4 | 0.264 | 1.69E-12 | 2.85E-17 | 0.9863,1.009,1.0577 | -0.0109,0.0073,0.0444 |
| | 4 | rs10016841 | 20213781 | SLIT2 | 0.133 | 1.95E-09 | 2.11E-13 | 0.9898,1.0272,1.0621 | -0.007,0.0187,0.0468 |
| | 4 | rs12507026 | 45181334 | GNPDA2 | 0.434 | 1.84E-11 | 6.78E-41 | 0.9762,0.9998,1.0381 | -0.0243,-6e-04,0.0435 |

| | | | | | | | | | |
|---|---|---|---|---|---|---|---|---|---|
| | 6 | rs34817112 | 27176628 | PRSS16 | 0.134 | 8.48E-17 | 3.51E-08 | 0.9857,1.0416,1.0635 | -0.0054,0.0154,0.026 |
| | 6 | rs3132947 | 32176782 | GPSM3 | 0.218 | 2.36E-13 | 8.53E-15 | 0.9834,1.0214,1.0563 | -0.0096,0.0119,0.038 |
| | 6 | rs987237 | 50803050 | TFAP2B | 0.18 | 2.18E-16 | 7.51E-43 | 0.9842,1.0249,1.0845 | -0.0148,0.0247,0.0833 |
| | 8 | rs17150703 | 9745798 | MSRA | 0.104 | 1.38E-09 | 2.05E-11 | 0.9925,1.0243,1.1486 | -0.0053,0.0196,0.0583 |
| | 10 | rs4132670 | 114767771 | TCF7L2 | 0.312 | 3.88E-11 | 2.75E-15 | 1.0205,0.986,0.9606 | 0.0133,-0.0084,-0.0263 |
| | 11 | rs2049045 | 27694241 | BDNF | 0.187 | 6.91E-10 | 8.20E-42 | 1.0115,0.9794,0.9461 | 0.0162,-0.0288,-0.0563 |
| | 12 | rs7132908 | 50263148 | BCDIN3D | 0.385 | 3.73E-11 | 3.94E-32 | 0.9791,1.0024,1.0429 | -0.0211,0.0046,0.0392 |
| | 12 | rs11057413 | 124489162 | ZNF664-FAM101A | 0.334 | 1.05E-10 | 6.30E-09 | 0.981,1.0071,1.0456 | -0.0104,0.0064,0.0173 |
| | 16 | rs4072402 | 28937259 | RABEP2 | 0.337 | 5.55E-12 | 2.72E-28 | 0.9802,1.0076,1.0463 | -0.0185,0.0081,0.0393 |
| | 16 | rs12716979 | 31011821 | STX1B | 0.375 | 1.40E-16 | 7.30E-24 | 1.031,0.9897,0.9517 | 0.0186,-0.0053,-0.0328 |
| | 16 | rs11642015 | 53802494 | FTO | 0.404 | 1.73E-73 | 7.43E-217 | 0.9398,1.0013,1.1095 | -0.0555,0.005,0.1062 |
| | 18 | rs10871777 | 57851763 | MC4R | 0.236 | 1.73E-19 | 3.01E-81 | 0.9767,1.0232,1.0751 | -0.0248,0.0262,0.0897 |
| | 19 | rs2238691 | 46179043 | GIPR | 0.194 | 3.46E-15 | 2.31E-32 | 1.0176,0.9706,0.9309 | 0.0142,-0.0231,-0.0537 |
| WC | 1 | rs10913469 | 177913519 | SEC16B | 0.205 | 3.80E-14 | 4.50E-44 | 0.9848,1.0189,1.0695 | -0.0166,0.0229,0.0724 |
| | 2 | rs62104180 | 466003 | FAM150B | 0.05 | 3.93E-14 | 4.02E-44 | 1.0061,0.9417,0.8124 | 0.0077,-0.0689,-0.1472 |
| | 2 | rs13412194 | 653245 | TMEM18 | 0.172 | 9.76E-15 | 1.39E-55 | 1.0134,0.9726,0.9343 | 0.0176,-0.0341,-0.0761 |
| | 3 | rs7649970 | 12392272 | PPARG | 0.121 | 5.60E-10 | 5.30E-10 | 0.9915,1.0245,1.0873 | -0.0057,0.0186,0.0314 |
| | 4 | rs12507026 | 45181334 | GNPDA2 | 0.434 | 2.39E-11 | 9.40E-31 | 0.9757,1.0016,1.0355 | -0.0207,-8e-04,0.0377 |
| | 6 | rs13198716 | 26582035 | ABT1 | 0.109 | 4.89E-15 | 0.0305 | 0.99,1.0363,1.0714 | -0.002,0.0078,0.0046 |
| | 6 | rs1062070 | 32148031 | RNF5 | 0.199 | 7.20E-12 | 6.06E-10 | 0.9862,1.0221,1.043 | -0.0079,0.0132,0.0217 |
| | 6 | rs4472337 | 34769765 | UHRF1BP1 | 0.155 | 5.60E-11 | 1.78E-23 | 0.9893,1.0237,1.0573 | -0.0105,0.024,0.0493 |
| | 6 | rs987237 | 50803050 | TFAP2B | 0.18 | 5.43E-12 | 1.09E-34 | 0.9874,1.0199,1.0669 | -0.0137,0.0239,0.0663 |

| | | | | | | | | | |
|---|---|---|---|---|---|---|---|---|---|
| | 7 | rs12667251 | 130449458 | KLF14 | 0.436 | 6.82E-12 | 1.91E-05 | 1.025,0.9962,0.9637 | 0.0096,-0.0023,-0.0109 |
| | 12 | rs7133378 | 124409502 | CCDC92 | 0.318 | 6.25E-10 | 0.506 | 0.9845,1.0071,1.0384 | 0.001,1e-04,-0.0034 |
| | 16 | rs8056890 | 28897452 | ATP2A1 | 0.355 | 5.57E-15 | 8.85E-40 | 0.9759,1.009,1.0433 | -0.0234,0.0094,0.0433 |
| | 16 | rs34898535 | 31025641 | STX1B | 0.378 | 1.11E-11 | 6.24E-22 | 1.0246,0.991,0.9616 | 0.0173,-0.0047,-0.0316 |
| | 16 | rs1421085 | 53800954 | FTO | 0.404 | 3.27E-52 | 3.21E-166 | 0.9501,1.0048,1.0807 | -0.0481,0.0038,0.0936 |
| | 18 | rs11152213 | 57852948 | MC4R | 0.236 | 5.62E-15 | 1.39E-70 | 0.9828,1.0153,1.0646 | -0.0224,0.0224,0.0898 |
| | 19 | rs1800437 | 46181392 | GIPR | 0.194 | 2.05E-11 | 1.19E-24 | 1.0137,0.9791,0.93 | 0.0124,-0.0203,-0.0445 |
| HC | 1 | rs6685593 | 203516075 | OPTC | 0.495 | 5.99E-11 | 2.47E-12 | 0.9682,1.0032,1.0238 | -0.016,-3e-04,0.0181 |
| | 1 | rs2605098 | 219643649 | LYPLAL1 | 0.338 | 1.11E-20 | 3.87E-38 | 0.9723,1.0085,1.0687 | -0.0209,0.0081,0.0483 |
| | 2 | rs62104180 | 466003 | FAM150B | 0.05 | 1.58E-09 | 5.56E-45 | 1.0054,0.9447,0.9202 | 0.0078,-0.07,-0.1422 |
| | 2 | rs6751993 | 635864 | TMEM18 | 0.167 | 3.84E-12 | 1.43E-58 | 1.0142,0.9714,0.932 | 0.0186,-0.0337,-0.0883 |
| | 2 | rs10200566 | 25130462 | ADCY3 | 0.451 | 2.32E-10 | 2.29E-18 | 0.9811,0.9976,1.0342 | -0.0171,0,0.026 |
| | 6 | rs34158769 | 26336572 | BTN3A2 | 0.104 | 5.06E-15 | 4.20E-13 | 0.9881,1.0436,1.0925 | -0.0061,0.0238,0.0417 |
| | 6 | rs3132947 | 32176782 | GPSM3 | 0.218 | 5.34E-11 | 2.52E-24 | 0.9851,1.019,1.049 | -0.0132,0.0176,0.0429 |
| | 6 | rs72891717 | 50858235 | TFAP2B | 0.169 | 6.03E-10 | 1.10E-36 | 0.9869,1.0226,1.0786 | -0.0134,0.0247,0.0784 |
| | 6 | rs141783576 | 127439897 | RSPO3 | 0.067 | 1.57E-10 | 4.72E-32 | 1.0064,0.9482,0.994[b] | 0.0079,-0.0512,-0.0833 |
| | 7 | rs17789506 | 130445574 | KLF14 | 0.493 | 2.80E-11 | 1.87E-18 | 0.9676,1.0001,1.0321 | -0.0189,-0.0019,0.0234 |
| | 12 | rs10846580 | 124415453 | CCDC92 | 0.337 | 7.64E-12 | 9.27E-17 | 0.9793,1.0122,1.0302 | -0.016,0.0106,0.0209 |
| | 16 | rs8056890 | 28897452 | ATP2A1 | 0.355 | 1.27E-09 | 1.71E-41 | 0.9762,1.0105,1.0362 | -0.0239,0.0096,0.0444 |
| | 16 | rs34898535 | 31025641 | STX1B | 0.378 | 2.57E-12 | 1.58E-27 | 1.0248,0.9945,0.9502 | 0.0198,-0.0054,-0.0352 |
| | 16 | rs1421085 | 53800954 | FTO | 0.404 | 1.65E-48 | 2.05E-152 | 0.9486,1.0029,1.0909 | -0.0462,0.0039,0.0893 |
| | 18 | rs11152213 | 57852948 | MC4R | 0.236 | 2.39E-16 | 1.44E-72 | 0.98,1.0189,1.0704 | -0.0237,0.0257,0.0817 |
| | 19 | rs2238691 | 46179043 | GIPR | 0.194 | 4.52E-11 | 9.80E-20 | 1.0162,0.9727,0.9364 | 0.0105,-0.0164,-0.0467 |

| | | | | | | | | | |
|---|---|---|---|---|---|---|---|---|---|
| WHR | 5 | rs459193 | 55806751 | C5orf67 | 0.253 | 2.86E-13 | 1.75E-19 | 0.9859,1.0102,1.0584 | -0.0128,0.0129,0.0354 |
| BFP | 1 | rs2820468 | 219673705 | LYPLAL1 | 0.345 | 3.76E-11 | 3.46E-21 | 0.9824,1.0063,1.0364 | -0.0164,0.0066,0.0328 |
| | 2 | rs1128249 | 165528624 | GRB14 | 0.392 | 1.93E-09 | 2.32E-18 | 0.9819,1.0045,1.0279 | -0.0165,0.0039,0.0275 |
| | 3 | rs900399 | 156798732 | CCNL1 | 0.397 | 1.82E-09 | 0.000121 | 1.0198,0.9932,0.9746 | 0.0065,-4e-04,-0.0138 |
| | 6 | rs2523625 | 31315648 | HLA-B | 0.331 | 2.69E-10 | 0.0215 | 0.9852,1.0071,1.0331 | -0.0049,0.0039,0.0041 |
| | 16 | rs62033406 | 53824226 | FTO | 0.411 | 2.15E-11 | 1.44E-91 | 0.9806,0.9991,1.0359 | -0.0367,0.0025,0.0677 |
| BMR | 6 | rs10456362 | 28221816 | ZKSCAN4 | 0.161 | 1.48E-09 | 1.63E-09 | 0.99,1.0252,1.0064 | -0.0069,0.0163,0.0186 |
| | 11 | rs80083564 | 27733143 | BDNF | 0.136 | 1.15E-09 | 2.32E-22 | 0.9895,1.0273,1.0718 | -0.0092,0.0242,0.0657 |
| | 12 | rs78719460 | 133395038 | GOLGA3 | 0.31 | 1.66E-09 | 1.92E-12 | 0.984,1.0063,1.0467 | -0.0108,0.0054,0.0289 |
| | 16 | rs1421085 | 53800954 | FTO | 0.404 | 8.95E-47 | 9.23E-154 | 0.9561,1.0008,1.0805 | -0.0469,0.0041,0.09 |
| | 18 | rs476828 | 57852587 | MC4R | 0.237 | 1.87E-20 | 1.35E-148 | 0.9775,1.0195,1.0716 | -0.0351,0.0388,0.1125 |

Note: a) p values smaller than $2.0\times10^{-9}$ are highlighted in pink; b) vQTLs with non-additive genetic effect on variance are highlighted in yellow.

**Supplementary Table 3. Environmental data used in the GEI analyses in the UKB**

| Item | Description | UDI |
|---|---|---|
| Sex | Sex | 31-0.0 |
| Age | Year of birth | 34-0.0 |
| DayW | Number of days/week walked 10+ minutes | 864-0.0 |
| DurW | Duration of walks | 874-0.0 |
| DayM | Number of days/week of moderate physical activity 10+ minutes | 884-0.0 |
| DurM | Duration of moderate activity | 894-0.0 |
| DayV | Number of days/week of vigorous physical activity 10+ minutes | 904-0.0 |
| DurV | Duration of vigorous activity | 914-0.0 |
| TimeD | Time spent driving | 1090-0.0 |
| TimeC | Time spent using computer | 1080-0.0 |
| TimeTV | Time spent watching television (TV) | 1070-0.0 |
| CurS | Current tobacco smoking | 1239-0.0 |
| PastS | Past tobacco smoking | 1249-0.0 |

Note: UDI, the Unique Data Identifier in the UKB dataset.

**Supplementary Table 4. GEI analyses between the 75 vQTLs and five environmental factors in the UKB**

| Trait | CHR | SNP | BP | Nearest Gene | P values of GEI analyses with | | | | |
|---|---|---|---|---|---|---|---|---|---|
| | | | | | Sex | Age | PA | SB | Smoking |
| FFR | 4 | rs6537292 | 145469968 | HHIP | 3.13E-02 | 7.20E-01 | 5.91E-01 | 6.39E-01 | 8.14E-02 |
| | 5 | rs12374521 | 147836880 | FBXO38 | 9.46E-02 | 1.43E-01 | 2.63E-01 | 4.79E-02 | 2.88E-04 |
| | 15 | rs56077333 | 78899003 | CHRNA3 | 2.36E-02 | 2.52E-02 | 9.88E-01 | 3.02E-04 | 4.55E-25 |
| BMD | 1 | rs1414660 | 240586695 | GREM2 | 7.60E-01 | 7.09E-05 | 1.45E-01 | 4.26E-01 | 4.55E-01 |
| | 6 | rs9371221 | 151885986 | CCDC170 | 9.81E-01 | 7.05E-01 | 8.33E-01 | 7.69E-01 | 1.28E-01 |
| | 6 | rs3020332 | 152008924 | ESR1 | 2.08E-01 | 4.76E-01 | 3.28E-01 | 9.46E-01 | 9.53E-01 |
| | 7 | rs4576334 | 38153747 | STARD3NL | 2.85E-01 | 9.08E-02 | 1.17E-01 | 4.58E-01 | 4.45E-01 |
| | 7 | rs10254825 | 120956440 | WNT16 | 3.06E-04 | 1.16E-07 | 4.02E-01 | 7.83E-01 | 1.59E-03 |
| | 11 | rs603140 | 86884615 | TMEM135 | 1.07E-03 | 2.75E-02 | 5.74E-01 | 4.58E-02 | 7.51E-01 |
| BW | 3 | rs13322435 | 156795468 | CCNL1 | 8.46E-02 | 1.44E-01 | 6.87E-01 | 3.69E-01 | 9.36E-01 |
| BMI | 1 | rs545608 | 177899121 | SEC16B | 8.59E-03 | 1.24E-04 | 6.11E-03 | 1.27E-02 | 7.51E-01 |
| | 1 | rs6689335 | 219628682 | LYPLAL1 | 6.65E-01 | 1.90E-01 | 3.15E-02 | 1.13E-01 | 7.38E-01 |
| | 2 | rs62104180 | 466003 | FAM150B | 2.75E-01 | 5.36E-02 | 2.52E-04 | 2.07E-01 | 1.48E-01 |
| | 2 | rs6751993 | 635864 | TMEM18 | 7.08E-01 | 1.01E-07 | 1.49E-02 | 9.67E-01 | 2.06E-01 |
| | 2 | rs10203386 | 25136866 | ADCY3 | 2.52E-01 | 4.94E-02 | 8.81E-03 | 8.92E-01 | 4.51E-01 |
| | 2 | rs1641155 | 58965211 | FANCL | 8.94E-01 | 4.64E-01 | 9.82E-01 | 3.09E-01 | 3.73E-01 |
| | 3 | rs1225053 | 131642852 | CPNE4 | 1.78E-01 | 7.44E-01 | 9.53E-01 | 8.74E-01 | 4.52E-01 |
| | 4 | rs10016841 | 20213781 | SLIT2 | 4.41E-01 | 8.52E-01 | 8.64E-03 | 5.42E-03 | 3.88E-03 |
| | 4 | rs12507026 | 45181334 | GNPDA2 | 1.59E-01 | 6.19E-02 | 6.40E-02 | 1.46E-01 | 5.21E-01 |
| | 6 | rs34817112 | 27176628 | PRSS16 | 9.93E-01 | 1.91E-02 | 1.52E-01 | 9.24E-03 | 9.21E-01 |

| | | | | | | | | | |
|---|---|---|---|---|---|---|---|---|---|
| | 6 | rs3132947 | 32176782 | GPSM3 | 2.18E-01 | 1.92E-03 | 6.16E-01 | 1.41E-01 | 3.66E-01 |
| | 6 | rs987237 | 50803050 | TFAP2B | 1.24E-01 | 1.41E-02 | 4.31E-01 | 1.68E-01 | 2.49E-02 |
| | 8 | rs17150703 | 9745798 | MSRA | 2.62E-01 | 7.88E-01 | 3.15E-01 | 6.25E-02 | 9.09E-01 |
| | 10 | rs4132670 | 114767771 | TCF7L2 | 3.03E-01 | 5.72E-01 | 1.73E-03 | 6.84E-04 | 2.36E-01 |
| | 11 | rs2049045 | 27694241 | BDNF | 1.67E-01 | 2.66E-01 | 1.59E-02 | 9.22E-01 | 2.62E-01 |
| | 12 | rs7132908 | 50263148 | BCDIN3D | 2.73E-01 | 2.94E-01 | 1.36E-03 | 2.15E-07 | 5.88E-04 |
| | 12 | rs11057413 | 124489162 | ZNF664-FAM101A | 1.02E-01 | 2.03E-01 | 6.54E-02 | 5.81E-03 | 9.57E-01 |
| | 16 | rs4072402 | 28937259 | RABEP2 | 9.25E-01 | 1.30E-01 | 1.82E-02 | 2.72E-03 | 3.58E-01 |
| | 16 | rs12716979 | 31011821 | STX1B | 6.74E-03 | 9.39E-01 | 7.48E-03 | 2.07E-01 | 5.89E-01 |
| | 16 | rs11642015 | 53802494 | FTO | 5.01E-03 | 2.35E-04 | 1.28E-10 | 1.64E-09 | 9.24E-05 |
| | 18 | rs10871777 | 57851763 | MC4R | 4.63E-01 | 3.72E-03 | 3.52E-04 | 1.41E-02 | 3.22E-02 |
| | 19 | rs2238691 | 46179043 | GIPR | 4.83E-01 | 7.04E-01 | 5.95E-02 | 1.74E-04 | 6.53E-01 |
| WC | 1 | rs10913469 | 177913519 | SEC16B | 1.63E-01 | 4.96E-03 | 2.74E-02 | 1.31E-01 | 2.15E-01 |
| | 2 | rs62104180 | 466003 | FAM150B | 6.08E-02 | 1.46E-01 | 1.04E-04 | 4.77E-01 | 5.19E-01 |
| | 2 | rs13412194 | 653245 | TMEM18 | 4.87E-01 | 1.88E-07 | 3.70E-02 | 7.16E-01 | 3.15E-01 |
| | 3 | rs7649970 | 12392272 | PPARG | 6.35E-01 | 1.49E-01 | 8.87E-02 | 7.91E-01 | 9.55E-01 |
| | 4 | rs12507026 | 45181334 | GNPDA2 | 2.12E-01 | 2.10E-02 | 4.08E-01 | 6.70E-02 | 8.13E-01 |
| | 6 | rs13198716 | 26582035 | ABT1 | 1.52E-01 | 1.50E-04 | 3.11E-02 | 2.45E-03 | 5.51E-01 |
| | 6 | rs1062070 | 32148031 | RNF5 | 2.30E-01 | 3.46E-05 | 2.32E-01 | 3.13E-01 | 1.96E-01 |
| | 6 | rs4472337 | 34769765 | UHRF1BP1 | 2.03E-01 | 6.15E-01 | 3.14E-02 | 5.56E-03 | 4.73E-01 |
| | 6 | rs987237 | 50803050 | TFAP2B | 2.52E-02 | 9.32E-03 | 5.61E-01 | 2.19E-01 | 3.32E-02 |
| | 7 | rs12667251 | 130449458 | KLF14 | 2.61E-01 | 6.30E-01 | 6.99E-01 | 3.50E-01 | 8.05E-01 |
| | 12 | rs7133378 | 124409502 | CCDC92 | 3.59E-03 | 1.45E-01 | 7.05E-01 | 3.62E-03 | 6.79E-01 |

| | | | | | | | | | |
|---|---|---|---|---|---|---|---|---|---|
| | 16 | rs8056890 | 28897452 | ATP2A1 | 8.75E-01 | 1.52E-01 | 3.93E-02 | 4.24E-03 | 2.10E-01 |
| | 16 | rs34898535 | 31025641 | STX1B | 5.92E-03 | 7.08E-01 | 1.69E-02 | 2.92E-01 | 9.87E-01 |
| | 16 | rs1421085 | 53800954 | FTO | 3.04E-02 | 2.17E-04 | 1.44E-07 | 2.84E-08 | 1.10E-02 |
| | 18 | rs11152213 | 57852948 | MC4R | 2.80E-02 | 1.36E-02 | 2.21E-03 | 3.39E-02 | 2.14E-01 |
| | 19 | rs1800437 | 46181392 | GIPR | 1.84E-01 | 4.59E-01 | 1.62E-01 | 9.50E-04 | 1.81E-01 |
| HC | 1 | rs6685593 | 203516075 | OPTC | 6.22E-01 | 8.84E-01 | 3.48E-01 | 7.42E-02 | 3.80E-01 |
| | 1 | rs2605098 | 219643649 | LYPLAL1 | 8.00E-03 | 8.14E-01 | 5.37E-02 | 4.52E-01 | 9.48E-01 |
| | 2 | rs62104180 | 466003 | FAM150B | 3.95E-01 | 2.89E-01 | 2.27E-05 | 4.70E-02 | 1.62E-01 |
| | 2 | rs6751993 | 635864 | TMEM18 | 6.50E-01 | 2.35E-04 | 1.87E-02 | 5.09E-01 | 3.11E-01 |
| | 2 | rs10200566 | 25130462 | ADCY3 | 2.35E-01 | 1.73E-01 | 3.57E-02 | 7.90E-01 | 7.74E-01 |
| | 6 | rs34158769 | 26336572 | BTN3A2 | 4.31E-01 | 1.05E-02 | 1.14E-02 | 1.60E-02 | 9.93E-01 |
| | 6 | rs3132947 | 32176782 | GPSM3 | 9.79E-01 | 4.23E-03 | 2.53E-01 | 3.71E-01 | 7.46E-02 |
| | 6 | rs72891717 | 50858235 | TFAP2B | 2.32E-01 | 2.19E-02 | 5.21E-01 | 8.67E-01 | 1.67E-01 |
| | 6 | rs141783576 | 127439897 | RSPO3 | 5.75E-01 | 5.59E-01 | 5.95E-02 | 8.08E-02 | 6.84E-01 |
| | 7 | rs17789506 | 130445574 | KLF14 | 6.09E-05 | 4.28E-01 | 5.89E-02 | 1.36E-01 | 2.73E-01 |
| | 12 | rs10846580 | 124415453 | CCDC92 | 6.97E-02 | 1.10E-01 | 4.89E-01 | 1.93E-02 | 6.48E-01 |
| | 16 | rs8056890 | 28897452 | ATP2A1 | 7.53E-01 | 2.66E-01 | 1.68E-02 | 2.05E-02 | 5.89E-01 |
| | 16 | rs34898535 | 31025641 | STX1B | 1.60E-02 | 4.20E-01 | 1.62E-02 | 1.94E-01 | 2.77E-01 |
| | 16 | rs1421085 | 53800954 | FTO | 1.18E-01 | 2.05E-03 | 5.32E-07 | 2.17E-06 | 1.94E-04 |
| | 18 | rs11152213 | 57852948 | MC4R | 2.38E-01 | 2.19E-02 | 9.80E-04 | 1.41E-02 | 6.68E-03 |
| | 19 | rs2238691 | 46179043 | GIPR | 8.71E-02 | 9.12E-01 | 2.66E-01 | 2.64E-04 | 2.85E-01 |
| WHR | 5 | rs459193 | 55806751 | C5orf67 | 9.25E-02 | 4.82E-01 | 1.92E-01 | 3.48E-04 | 6.67E-01 |
| BFP | 1 | rs2820468 | 219673705 | LYPLAL1 | 8.76E-01 | 1.19E-01 | 1.60E-02 | 8.66E-02 | 2.40E-01 |

| | | | | | | | | | |
|---|---|---|---|---|---|---|---|---|---|
| | 2 | rs1128249 | 165528624 | GRB14 | 3.18E-01 | 2.91E-02 | 3.55E-01 | 3.41E-04 | 5.58E-01 |
| | 3 | rs900399 | 156798732 | CCNL1 | 1.04E-05 | 2.63E-01 | 2.15E-01 | 1.13E-02 | 1.64E-02 |
| | 6 | rs2523625 | 31315648 | HLA-B | 2.66E-01 | 2.87E-01 | 1.35E-01 | 6.76E-01 | 6.38E-01 |
| | 16 | rs62033406 | 53824226 | FTO | 1.14E-02 | 2.47E-04 | 4.43E-03 | 3.52E-02 | 1.02E-01 |
| BMR | 6 | rs10456362 | 28221816 | ZKSCAN4 | 8.90E-01 | 8.38E-03 | 6.68E-01 | 1.60E-01 | 9.13E-01 |
| | 11 | rs80083564 | 27733143 | BDNF | 4.20E-01 | 1.69E-02 | 1.00E-01 | 5.22E-01 | 9.89E-01 |
| | 12 | rs78719460 | 133395038 | GOLGA3 | 3.41E-01 | 8.69E-01 | 3.04E-01 | 9.40E-01 | 6.67E-01 |
| | 16 | rs1421085 | 53800954 | FTO | 2.07E-01 | 2.60E-07 | 1.52E-06 | 1.45E-07 | 1.54E-03 |
| | 18 | rs476828 | 57852587 | MC4R | 1.20E-01 | 1.22E-03 | 9.98E-03 | 6.47E-02 | 1.62E-02 |

Note: p values smaller than $1.3\times10^{-4}$ are highlighted in pink.

**Supplementary Table 5. GEI effect between the *CHRNA5-A3-B4* locus and smoking on FFR**

| Phenotype | Top vQTL SNP | Effect size (Standard error) | | P values | | |
|---|---|---|---|---|---|---|
| | | never smokers (n= 188,860) | ever smokers (n= 160,488) | vQTL analysis | QTL analysis | GEI test |
| FFR | rs56077333 | 0.0105 (0.0035) | -0.0453 (0.0042) | 1.09E-14 | 2.11E-06 | 4.55E-25 |

**Supplementary Table 6. GEI effect between the *WNT16-CPED1* locus and age on BMD**

| Phenotype | Top vQTL SNP | Effect size (Standard error) | | | | *P* values | | |
|---|---|---|---|---|---|---|---|---|
| | | Age group 1: 40-49 years ($n$ = 59,734) | Age group 2: 50-59 years ($n$ = 108,736) | Age group 3: 60-69 years ($n$ = 156,173) | Age group 4: 70-74 years ($n$ = 23,250) | vQTL analysis | QTL analysis | GEI test |
| BMD | rs10254825 | 0.1448 (0.0081) | 0.1650 (0.0059) | 0.1907 (0.0050) | 0.1765 (0.0119) | 2.01E-45 | 0 | 1.16E-07 |

**Supplementary Table 7. Associations of *FTO* locus with obesity-related traits stratified by physical activity (PA) levels**

| Phenotype | Top vQTL SNP | Effect size (Standard error) | | | *P* values | | |
|---|---|---|---|---|---|---|---|
| | | Low PA group ($n$ = 103,374) | Intermediate PA group ($n$ = 145,889) | High PA group ($n$ = 97,506) | vQTL analysis | QTL analysis | GEI test |
| BMI | rs11642015 | 0.1018 (0.0049) | 0.0715 (0.0037) | 0.0609 (0.0041) | 1.73E-73 | 7.43E-217 | 1.28E-10 |
| WC | rs1421085 | 0.0858 (0.0048) | 0.0652 (0.0037) | 0.0524 (0.0042) | 3.27E-52 | 3.21E-166 | 1.44E-07 |
| HC | rs1421085 | 0.0825 (0.0049) | 0.0623 (0.0037) | 0.0505 (0.0042) | 1.65E-48 | 2.05E-152 | 5.32E-07 |
| BMR | rs1421085 | 0.0842 (0.0049) | 0.0608 (0.0037) | 0.0531 (0.0043) | 8.95E-47 | 9.23E-154 | 1.52E-06 |

**Supplementary Table 8. Associations of *FTO* locus with obesity-related traits stratified by sedentary behaviour (SB) levels**

| | | Effect size (Standard error) | | | *P* values | | |
|---|---|---|---|---|---|---|---|
| Phenotype | Top vQTL SNP | SB group 1: 0-5 hours (*n* = 244,215) | SB group 2: 6-11 hours (*n* = 89,712) | SB group 3: 12-17 hours (*n* = 5,445) | vQTL analysis | QTL analysis | GEI test |
| BMI | rs11642015 | 0.0694 (0.0027) | 0.1001 (0.0052) | 0.1085 (0.0234) | 1.73E-73 | 7.43E-217 | 1.64E-09 |
| WC | rs1421085 | 0.0593 (0.0028) | 0.0879 (0.0050) | 0.1089 (0.0223) | 3.27E-52 | 3.21E-166 | 2.84E-08 |
| HC | rs1421085 | 0.0577 (0.0028) | 0.0815 (0.0052) | 0.1199 (0.0230) | 1.65E-48 | 2.05E-152 | 2.17E-06 |
| BMR | rs1421085 | 0.0576 (0.0028) | 0.0849 (0.0052) | 0.1024 (0.0233) | 8.95E-47 | 9.23E-154 | 1.45E-07 |
